# Structure and functional diversity of antibodies targeting the *P. falciparum* circumsporozoite protein C-terminal domain

**DOI:** 10.64898/2026.06.06.730512

**Authors:** Re’em Moskovitz, Iszac Burton, Nathan Beutler, Gonzalo Gonzalez-Paez, Thea Zalunardo, Michael V Bick, Justin Ndihokubwayo, Kiara Gambuzza, Jerry Zhao, Robyn L Stanfield, Xueyong Zhu, Monika Jain, Elizabeth A Winzeler, Daniel E Emerling, Christian F Ockenhouse, Randall S MacGill, Emily Locke, C Richter King, Dennis R Burton, Thomas F Rogers, Lars Hangartner, Ian A Wilson

## Abstract

The *Plasmodium falciparum* circumsporozoite protein (*Pf*CSP) is the major surface antigen on *Pf* sporozoites. WHO-recommended vaccines RTS,S/AS01_E_ and R21/Matrix-M target the *Pf*CSP major repeat region and C-terminal domain (ctCSP). Although multiple studies associated protection with antibody responses to ctCSP, only a few ctCSP-specific monoclonal antibodies (mAbs) have been characterized. Here, crystal structures of 11 Fab-ctCSP complexes reveal how mAbs against the conserved β-epitope region achieve diverse modes of strain-transcending recognition, in contrast to mAbs to the hypervariable ⍺-epitope. Consistent with previous studies, ctCSP on sporozoites could be unmasked by mAbs that bind CSP repeats, with unmasking dependent on the mAb fine-specificity and binding mode. In vitro, ctCSP mAbs promoted stronger Fc-receptor signaling, cellular cytotoxicity, and phagocytosis than repeat region mAbs, while mAb combinations targeting distinct *Pf*CSP epitopes modulated Fc-signaling and cellular cytotoxicity. This study provides a rationale for optimization of *Pf*CSP-based immunogens to enhance Fc-mediated contributions to malaria vaccine efficacy.

## Introduction

Despite global eradication efforts, malaria remains a significant global health burden, causing an estimated 282 million cases and 610 thousand deaths in 2024^1^. *Plasmodium falciparum* is responsible for over 90% of malaria cases in the World Health Organization (WHO) African region, and over 95% of malaria-associated deaths worldwide^1^. While global malaria prevention efforts have resulted in a decrease in malaria cases and deaths over the last two decades, this trend has stalled in recent years^1^. The success of early antimalarial therapies is jeopardized by the acquisition of antimalarial resistance and failure of front-line antimalarial therapies^2,3^, reinforcing the need for efficacious strain-transcending interventions.

*Plasmodium* sporozoites, the liver-infective form of the parasite, are deposited in the skin during the blood meal of an infected female *Anopheles* mosquito^4^. Sporozoites migrate out of the dermis through blood or lymph vessels towards the liver, invading hepatocytes to establish liver infection^4^. In the 1980s, the circumsporozoite protein (CSP) was identified as the immunodominant major surface antigen of sporozoites^5–7^, opening the door for recombinant CSP-based vaccination^8,9^. *Plasmodium* CSPs contain a conserved domain architecture: an N-terminal domain, a conserved junctional region (RI), a central repeat region, a C-terminal linker, and a C-terminal thrombospondin type-I (⍺TSR) domain, linked to the parasite membrane by a glycophosphatidylinositol (GPI) anchor^10,11^. The central repeat region of *P. falciparum* (3D7 strain) CSP contains 1 junctional NPDP repeat, 4 minor NVDP repeats, and ∼37 major NANP tetrapeptide repeats. The excellent immunogenicity of the NANP major repeats, improved by inclusion of T-cell epitopes found in the ⍺TSR domain, resulted in the development of RTS,S/AS01_E_ (Mosquirix^TM^). This first-generation subunit vaccine contains 19 NANP repeats and the CSP C-terminal domain (ctCSP) fused to the hepatitis B surface antigen (HBsAg) assembled into a virus-like particle (VLP) with unmodified HBsAg^12–15^. R21/Matrix-M contains an identical CSP-HBsAg component as RTS,S/AS01_E_, but is assembled without unmodified HBsAg to present CSP at a higher density. When administered seasonally to children and infants in regions of high malaria transmission, both vaccines reduce malaria cases by around 65–75%^16–21^. However, protection provided by both vaccines is below the WHO goal of 90%, vaccine efficacies are inconsistent across age groups, and wane rapidly following the final immunization^22–25^. These factors highlight the need for improved understanding of vaccine-induced protection for development of next-generation immunogens.

In RTS,S/AS01_E_ and R21/Matrix-M vaccination, antibody responses towards the NANP major repeats have been associated with protection^20,22,26–30^, and potent monoclonal antibodies (mAbs) have been isolated from RTS,S vaccinees^31–34^. While all vaccines in development targeting the pre-erythrocytic stage of malaria contain components of the central repeat region^35^, some analyses of patient sera from recent RTS,S/AS01_E_ and R21/Matrix-M clinical trials did not identify the antibody response to the CSP repeats as a correlate of protection^36,37^. Other studies of RTS,S vaccinee sera, however, have associated ctCSP responses with protection independently of anti-NANP responses^30,36–41^. Nevertheless, there have been fewer studies on ctCSP binding mAbs^42–45^. The antibody response to ctCSP is dominated by V_H_3-21/V_L_3-21 lineage mAbs recognizing the ⍺-epitope on the polymorphic face of ctCSP, which limits binding and protection to specific strains^33,42,44,45^. mAb 1512, isolated from a participant in the MAL071 Phase 2a controlled human malaria infection (CHMI) study, binds the β-epitope on the conserved face of ctCSP, provides strain-transcending binding, and modest protection from sporozoite challenge in mice^33,36,44^. The frequency, breadth, and binding modes of antibodies towards the conserved β-epitope are largely unexplored. To characterize the antigenic landscape of ctCSP antibodies elicited by RTS,S/AS01_E_ vaccination^36^, we structurally characterized 11 ctCSP-binding mAbs from the MAL071 trial^33,36^, and identified 7 strain-transcending mAbs of diverse lineages binding the conserved β-epitope region as well as another conserved ctCSP epitope.

Interrogating the protective contributions of the antibody response to ctCSP is impeded by ctCSP masking on the surface of salivary gland sporozoites^46^. CtCSP is only naturally exposed during hepatocyte invasion, or by binding of some antibodies to the central repeat region^43,46,47^. We used our antibody toolkit to reveal the impact of repeat antibody fine specificity on ctCSP unmasking and antibody competition on the sporozoite surface. Although our results indicate that our panel of antibodies is weakly protective in mice, we used *in vitro* models of human FcɣRIIIa signaling, neutrophil-mediated antibody-dependent cellular phagocytosis (ACDP), and natural killer (NK) cell-mediated antibody-dependent cellular cytotoxicity (ADCC) to reveal functional differences between antibodies targeting different regions of *Pf*CSP. The dynamics of additivity and inhibition in Fc-receptor signaling, ADCC, and ADCP produced by combining mAbs to multiple CSP epitopes provides molecular context for serological observations of Fc-mediated protection in RTS,S vaccinees. Together, our findings provide rationale for the optimization of next-generation CSP-based immunogens to improve vaccine efficacy by enhancing Fc-mediated contributions to protection.

## Results

The antibodies analyzed in this study were isolated from the MAL071 Phase 2a CHMI clinical trial^33,36^. Previously, paired heavy and light chain sequences were obtained from circulating peripheral blood mononuclear cells (PBMCs) isolated from vaccinees 7 days after the third immunization^33^, and a screening library of 369 mAb sequences was chosen representing dominant and unique lineages from both protected and unprotected vaccinees^33^. In addition to 8 previously characterized mAbs^44^, 12 mAbs bound ctCSP (residues 283-375 of PfCSP 3D7) by ELISA. V_H_ and V_L_ sequences of all ctCSP mAbs were subcloned into IgG1 heavy and light chain vectors, respectively, for recombinant expression of IgG and antigen binding fragments (Fab). The antibody sequence information is summarized in Table S1.

### MAL071 mAbs bind two dominant epitopes on the surface of *Pf*CSP ⍺TSR

First, we used bio-layer interferometry (BLI) to determine the competition groups of 14 antibody IgGs against ctCSP (Fig. 1A). Biotinylated ⍺TSR domain was immobilized on a streptavidin-coated biosensor surface, saturated with mAb1, and mAb2 binding amplitude was measured and normalized. Two predominant competition bins were established, the first containing previously characterized ⍺-epitope mAbs 234, 236, 352, 1488^44^ together with mAbs 1392, 1502, and 1525, and the second including β-epitope binding mAb 1512^44^ together with mAbs 369, 389, 1393, 1437, 1550 (Fig. 1A). In previous experiments, epitope binning suggested unilateral mAb competition whereby mAb binding to the ⍺-epitope abrogates binding to the β-epitope by an unknown mechanism, but mAb binding to the β-epitope did not abrogate binding to the ⍺-epitope ^44^. Our results did not show competition between ⍺- and β-epitope bins. A unique competition profile was only observed for mAb 1504 that produced a third epitope bin. We observed no competition for mAb binding (mAb2) to ⍺TSR domain previously saturated by 1504 (mAb1), but strong competition (<0.61 in Fig. 1A) for 1504 binding to ⍺TSR domain saturated with most other ctCSP mAbs (except mAbs 1525, 369, and 389).

**Figure 1.**
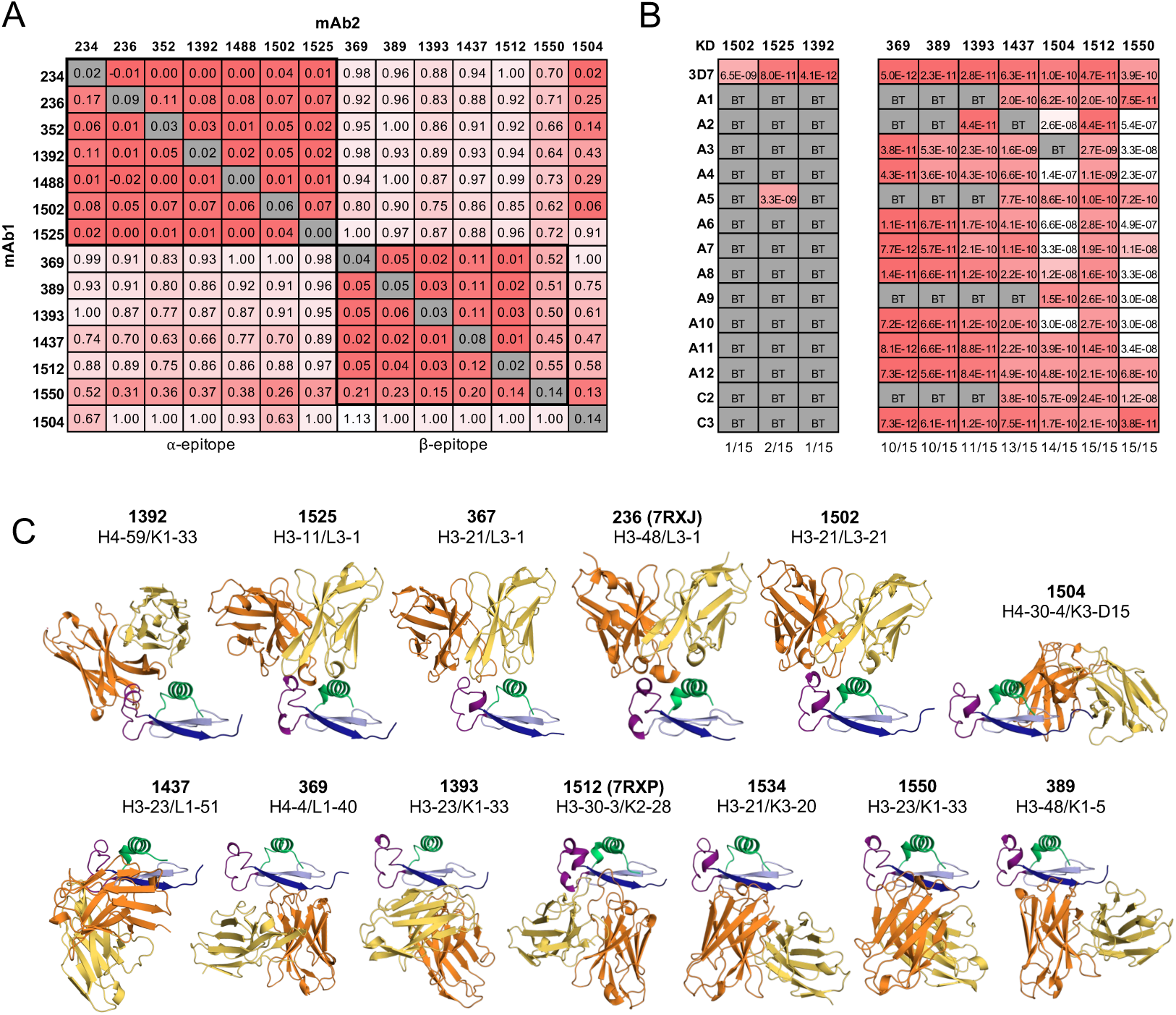
MAL071 mAbs bind two dominant ctCSP epitope regions. A) Epitope binning of MAL071 IgG binding to biotinylated ⍺TSR domain by bio-layer interferometry. B) Binding of MAL071 IgG to 15 ctCSP haplotypes including the 3D7 vaccine strain by surface plasmon resonance. BT = below threshold. Fraction of bound haplotypes shown at base of column. C) Overview of X-ray crystal structures of 11 new Fab-⍺TSR complexes, with Fab 1512 and Fab 236 included for reference. Structures are arranged by the relative disposition of the Fabs on the ⍺TSR domain, with ⍺-epitope and RII+ epitope Fabs shown on top and β-epitopes below. Fab heavy chain (orange), light chain (yellow), ⍺TSR Th2R (green), Th3R (purple), TSR-homology domain RII^+^ (light blue) and CS-T3 (dark blue) are shown in cartoon representation.

Next, we determined the binding breadth of MAL071 IgG against a panel of 15 recombinant ⍺TSR domain haplotypes by surface plasmon resonance (SPR) (Fig. 1B). The haplotypes chosen, including vaccine strain 3D7, represent genetic diversity detected across East and West Africa^25^ (listed in Beutler & Pholcharee et al.^44^). All ctCSP mAbs bound the vaccine strain 3D7 ⍺TSR domain with affinities (K_D_) in the nanomolar to picomolar range. Like other ⍺-epitope region specific mAbs, mAbs 1392, 1502, and 1525 demonstrated strain specificity^44^. In contrast, mAbs belonging to the β-epitope bin and mAb 1504 bound most haplotypes, with mAbs 1512 and 1550 binding all 15 haplotypes. Together, our epitope binning and breadth measurements confirmed that multiple MAL071 mAbs share an overlapping epitope with mAb 1512 and provide strain-transcending binding.

To further characterize fine epitopes of ctCSP mAbs, we determined crystal structures of 11 ctCSP Fabs in complex with a recombinant ⍺TSR domain (*Pf*CSP 310-375) at resolutions ranging from 1.6-2.3 Å (Fig. 1C). mAbs 367 and 1534 did not express in IgG format for epitope binning experiments but could be expressed in Fab format for crystallographic analysis. Crystallographic data collection and refinement statistics in Tables S2-4. Kabat numbering is used throughout for antibody sequences.

### mAbs bind to the polymorphic ⍺-epitope region with conserved interactions

Fabs 1392, 1502, 1525, and 367 bound the ⍺-epitope region (Fig. 1C) that was first defined by human mAb 1710 (isolated following immunization with radiation-attenuated sporozoites), and later by mAbs 234, 236, 352, and 1488 (isolated from RTS,S/AS01_E_ MAL071 vaccinees)^42,44^. Despite some relatively small variation in their angles of approach, all mAbs bound the ⍺-epitope region with a consistent footprint encompassing the Th2R ⍺-helix and Th3R CS-flap (Fig. 1C, 2A, and S1A) and produced 570-788 Å^2^ of buried surface area (BSA) on the ⍺TSR domain surface (Fig. S1A). As observed for other mAbs that targeted the ⍺-epitope region, V_H_3-21 Fab 367 and Fab 1502 and V_H_3-11 Fab 1525 use backbone and side chains of their Ser/Thr-rich HCDR2 to interact with the polymorphic Th3R Asp356 and Glu357 side chains (Fig. S2A and S2B). Similarly, their V_L_3-21 and V_L_3-1 light chains interact with the polymorphic Th2R Lys314, Lys317, and Asn321 (Fig. S2A and S2C). In all three Fabs, a hydrophobic HCDR3 residue (Ile, Ile, Leu) is inserted into the hydrophobic ⍺TSR domain core (Fig. S2A and S2D). In contrast to other characterized Fabs, Fab 1392 interacted with the ⍺-epitope region using only its V_H_4-50 heavy chain, which alone produced 737 Å^2^ of BSA on the antibody (Fig. S1B). Fab 1392 contains a 19aa long HCDR3 that extends from the Ig domain as an elongated β-hairpin, inserting between the Th3R CS-flap and Th2R ⍺-helix (Fig. S2A). In the Fab1392-⍺TSR structure, HCDR2 Asn54^H^ and Arg58^H^ engage the polymorphic Glu357 and Asp356, and HCDR3 Asp97^H^ and Arg100d^H^ interact with the adjacent Lys355 and Asp359, which stabilize the bound CS-flap in two ⍺-helical turns (Fig. S2B). The polymorphic Th2R Asn321 and Gln324 side chains interact with HCDR3 Asp100^H^ and the backbone of Tyr99^H^ and Phe100a^H^ (Fig. S2C), positioning the hydrophobic Phe100a^H^ side chain in the hydrophobic ⍺TSR domain core (Fig. S2D).

**Figure 2.**
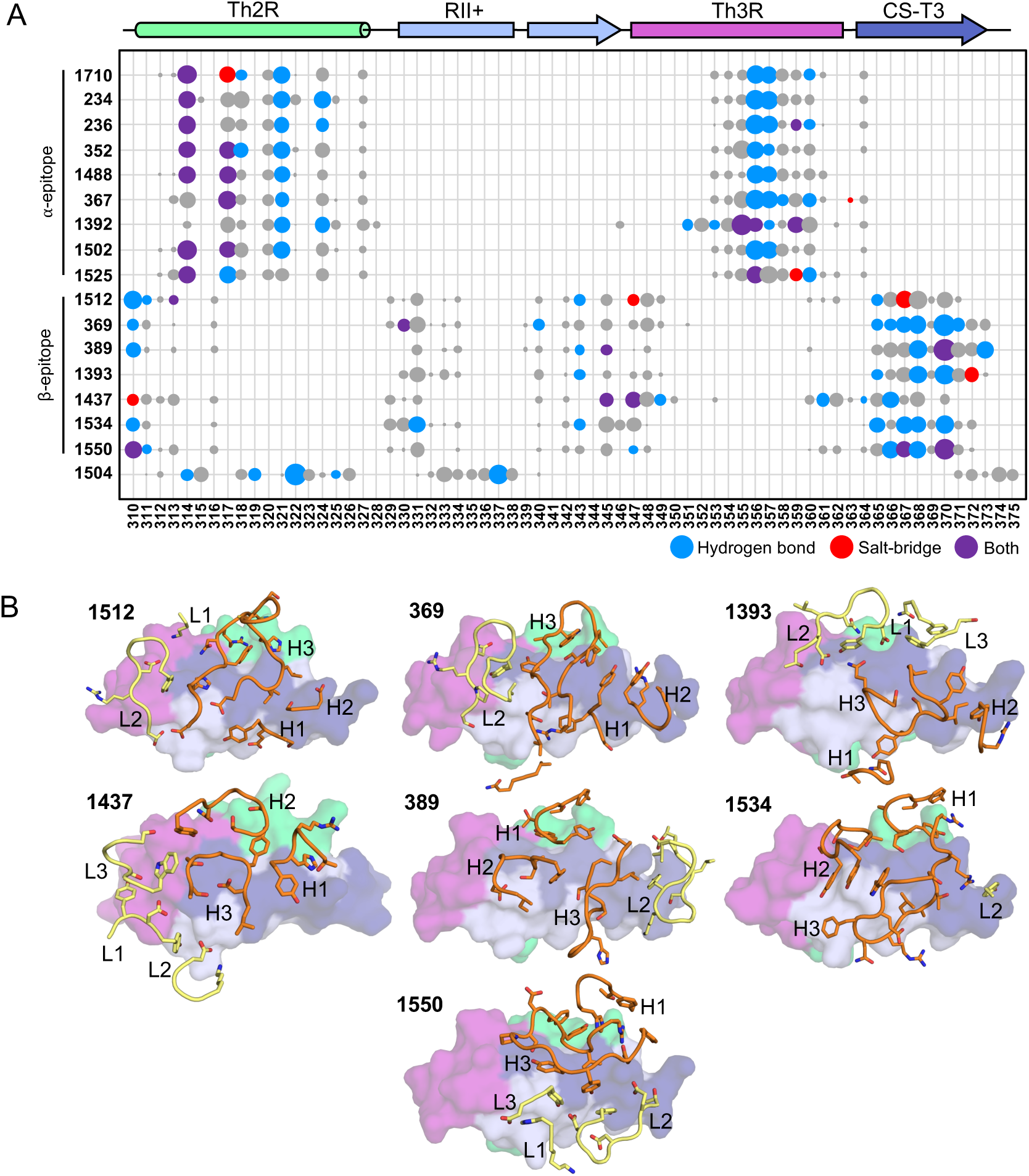
MAL071 mAbs bind the β-epitope region with diverse interactions and orientations. A) Bubble plot of per residue buried surface area (BSA) formed by Fab binding to the ⍺TSR surface surface with area of bubble proportional to BSA. Amino acids forming hydrogen bonds (blue), salt-bridge interactions (red), or both (purple) are mapped, highlighting binding features of ⍺TSR-binding Fabs. Previously published Fabs 1710 (PDB:6B0S), 234 (7RXI), 236 (7RXJ), 352 (7S0X), 1488 (7RXL), and 1512 (7RXP) are included for reference. B) Paratopes of β- epitope binding Fabs in complex with *Pf*CSP ⍺TSR domain. Fab heavy chain (orange) and light chain (yellow) backbone are shown in backbone tube representation with side chains of residues with >5Å^2^ BSA shown as sticks, and CDRs labeled H1–3 and L1–3. The ⍺TSR domain surface is shown with antibody binding to individual components of their epitope highlighted: Th2R in lime green, Th3R in magenta, RII^+^ in light blue, and CS-T3 in dark blue. Fab 1512 is shown for reference.

Across all mAbs that bound the ⍺-epitope region, the insertion of a hydrophobic HCDR3 side chain contributes over 10% of the BSA from a single residue, accommodating binding with diverse HCDR3 lengths and sequences by non-specific hydrophobic interactions (Tables S1, S5). By mapping the hydrogen-bond and salt-bridge interactions in the Fab-⍺TSR interfaces onto their BSA footprints, we show mAbs bind the ⍺-epitope region of the 3D7 ⍺TSR domain using specific polar and charged interactions with polymorphic residues Lys417, Lys317, Asn321, and Asn324 of the Th2R epitope, and Asp356 and Glu357 of the Th3R epitope, resulting in their observed strain specificity (Fig. 2A).

### The conserved β-epitope region is targeted by diverse lineages and binding modes

Fabs 369, 389, 1393, 1437, 1534, and 1550 bound the β-epitope region previously defined by Fab 1512 (Fig. 1C)^44^, approach the domain from a variety of angles, and produced 629-870 Å^2^ BSA on the ⍺TSR domain surface (Fig. S1A). Fabs 369, 1393, and 1437 bound the ⍺TSR domain in a roughly similar orientation as Fab 1512, with their heavy chain interacting with the C-terminal residues of the domain and light chain binding toward the CS-flap. This orientation positions HCDR3 to interact with the CS-T3 β-strand (Fig. 1 and 2B). Due to a rotated and more sideways approach, Fab 1437 forms more extensive contacts with the Th3R and Th2R epitopes (Fig. S1A). Fabs 389, 1534, and 1550 bind in the opposite orientation, interacting with the C-terminal residues of the domain using their LCDR2 while still maintaining binding with the CS-T3 β-strand through HCDR3 (Fig. 1 and 2B). Fab 1534 bound the ⍺TSR domain primarily through the heavy chain, which produced 741 Å^2^ of the total 761 Å^2^ BSA (Fig. S1A). With their varied orientations and V_H_/V_L_ pairing, β-epitope Fabs produce a footprint encompassing the N-terminal residues of both Th2R and CS-flap, with the majority of BSA produced by binding to the CS-T3 strand of the thrombospondin homology domain (Fig. 2A and S1A).

The residues composing the CS-T3 epitope provide a complete range of hydrophobic, aliphatic, acidic, and basic functional groups alongside exposed protein backbones as a platform for diverse binding modes across β-epitope binding mAbs. Ordered waters at the Fab:CS-T3 interface facilitate additional interactions, increasing contacts with the β-epitope region (Fig. S3). Mapping the hydrogen bonding and salt-bridge interactions onto their BSA footprint identifies that the majority of Fabs interact with Glu310, the backbone of Ile368, and Lys370, where the interactions are not as conserved compared to the ⍺-epitope region (Fig. 2A). Collectively, the broad recognition of ctCSP haplotypes by mAbs targeting the β-epitopes region provides a structural basis for the strain-transcending recognition of ctCSP and context for previous observations that associated such responses with protection in RTS,S vaccinee sera^39^.

### Fab 1504 binds a new conserved epitope on ctCSP

Fab 1504 bound the ⍺TSR domain with a distinct angle of approach (Fig. 1C). Primarily contacting the Th2R and RII^+^ regions, Fab 1504 defines a novel interaction interface on ctCSP (Fig. 1C and S4). With a 17aa HCDR3 in an extended β-hairpin conformation, the Fab 1504 heavy chain contacts the conserved facet of the Th2R helix with an interface abundant in hydrophobic and aromatic residues (Fig. S4A and S4B), which positions the light chain to interact with the CS-T3 epitope (Fig. S4C). The RII^+^ region is primarily engaged by the heavy chain, with Val97^H^ backbone forming a hydrogen bond to Thr337 hydroxyl that is conserved in the four Fab-⍺TSR complexes in the unit cell of our crystal structure (Fig. S4D and S4E). The RII^+^ sequence is highly conserved across *Plasmodium* species, thrombospondin domains, and other eukaryotic cell adhesion proteins^48,49^. Within this sequence, *Pf*CSP Thr337 is a conserved site for O-fucosylation^50^. Our ⍺TSR domain, however, was recombinantly expressed in *E. coli* lacking the appropriate O-fucosylation machinery. Alignment of O-fucosylated threonine onto Thr337 in our structure produces substantial spatial overlap with HCDR3 Val97^H^ and Asn100i^H^ (Fig. S4F) and would suggest that Fab 1504 could not bind O-fucosylated *Pf*CSP on the surface of sporozoites without substantial structural rearrangements.

### Binding of RTS,S-induced mAbs to ctCSP on the sporozoite surface requires ctCSP unmasking

On the surface of salivary gland sporozoites, ctCSP is masked by interaction with the N-terminal domain of CSP^47^, and some mAbs targeting the central repeats can unmask ctCSP by an unknown mechanism^43^. To investigate, we determined binding of MAL071 ctCSP mAbs to freshly isolated salivary gland *Plasmodium berghei* (*Pb*) sporozoites that express full-length *Pf*CSP (tg*Pb-Pf*CSP), in the presence or absence of a repeat-region mAb by flow cytometry. Sporozoites were incubated with 10 µg/ml ctCSP Alexa Fluor 568-labelled IgG and 10 µg/ml unlabeled 311 IgG, which binds the central repeat region of CSP^31,32,51^, or negative control Zika IgG^32^ (Fig. 3A, gating pathway in Fig. S5A). In the presence of 311 IgG, all ctCSP IgG bound over 90% of sporozoites, while in the absence of 311 IgG, most anti-ctCSP IgG did not (Fig. 3B). However, 369, 1437, 1525 IgG bound over 95% of sporozoites without 311 IgG, albeit with a ∼2-fold reduced median-fluorescent intensity (MFI). Unexpectedly, 1504 IgG bound over 90% of sporozoites and with a ∼4-fold higher MFI without 311 IgG (Fig. 3B). We also determined binding of a reduced panel of ctCSP IgG to transgenic *P. berghei* sporozoites in which the *Pb*CSP ⍺TSR is replaced with the *Pf*CSP ⍺TSR sequence, retaining the sequence of the *Pb*CSP N-terminal and central repeat domain (tg*Pb-Pf*ctCSP)^52^. Unexpectedly, the ⍺-epitope and β-epitope regions of the *Pf*CSP ⍺TSR domain were accessible for mAb binding on 90% of sporozoites in the absence of an antibody binding to the *Pb*CSP central-repeat, suggesting that masking of ctCSP may require interactions between the CSP N-terminal and C-terminal domains that are disrupted in tg*Pb-Pf*ctCSP parasites. In contrast, only 50% of sporozoites bound 1504 IgG, and with a 20-fold lower MFI (Fig. 3C). Furthermore, the smeared sporozoite population produced by 1504 binding may be indicative of rapid dissociation from sporozoites or heterogenous epitope presentation (Fig. S5B). Overall, these results show that MAL071 ctCSP mAbs can bind ctCSP on the surface of transgenic *Pb* sporozoites that express full length *Pf*CSP, and ctCSP unmasking is significantly enhanced by repeat mAb 311. While there does not appear to be a difference in accessibility to the ⍺- or β-epitope regions, CSP conformation or fucosylation may impact accessibility to the RII+ epitope.

**Figure 3.**
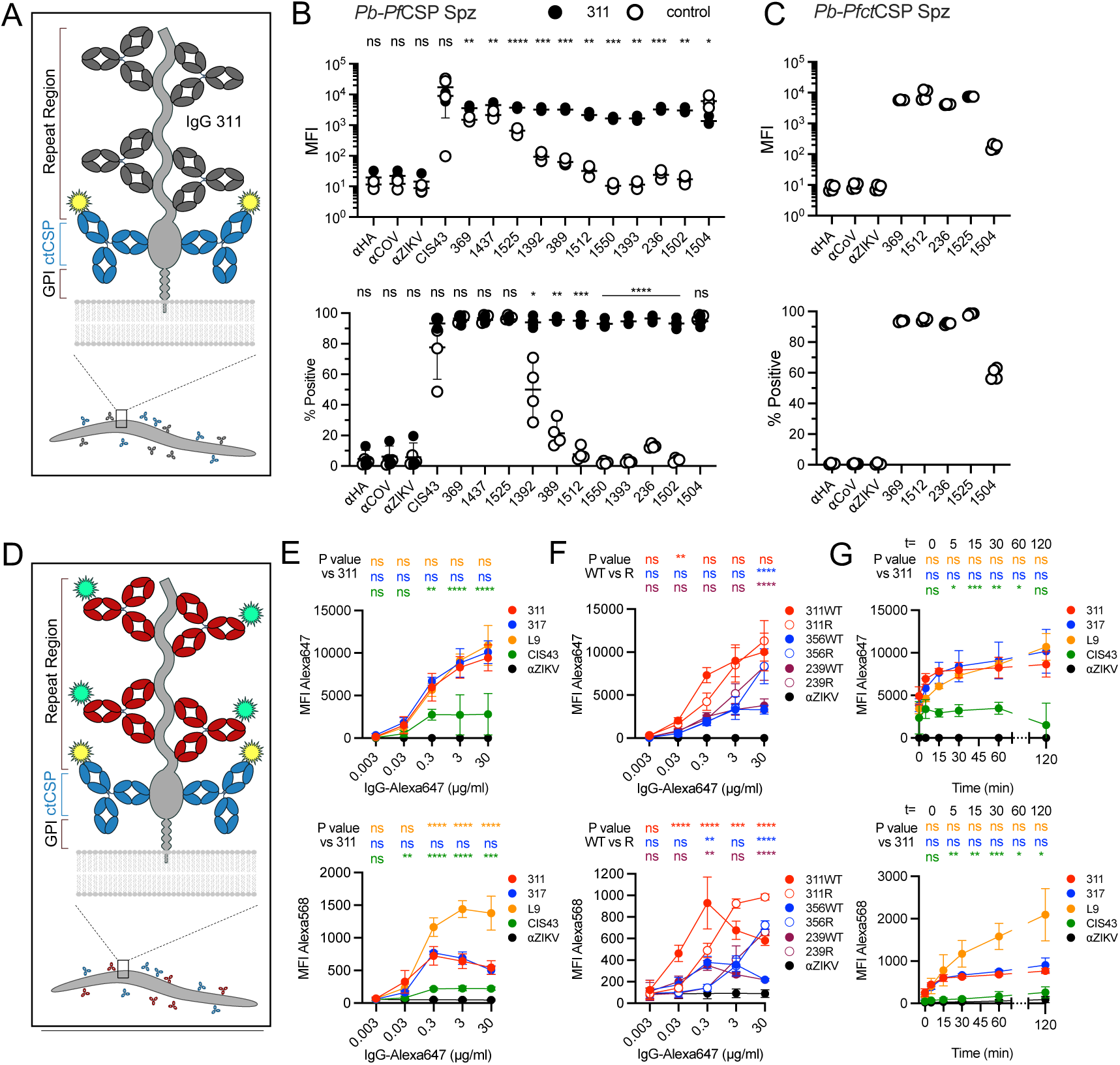
Effect of antibody specificity on ctCSP unmasking on sporozoites. A) Schematic of sporozoite staining to measure ctCSP mAb binding to *Pb-Pf*CSP sporozoites (Spz). B) Binding of Alexa Fluor 568-labelled mAbs to freshly isolated *Pb-Pf*CSP sporozoites in presence of mAb 311 or control IgG (N=4), including positive control CIS43 IgG, and negative control Influenza A hemagglutinin, SARS-CoV-2, and Zika IgGs. C) Binding of Alexa Fluor 568-labelled mAbs to freshly isolated *Pb-Pf*ctCSP sporozoites (N=2). D) Schematic of sporozoite staining to assess effect of repeat mAbs on ctCSP accessibility, with ctCSP mAbs in blue and repeat mAbs in red. E, F) Concentration-dependent binding of repeat-binding mAbs to *Pb-Pf*CSP sporozoites and associated mAb 1512-Alexa Fluor 568 binding (both N=3). G) Time-course of antibody binding and ctCSP unmasking on *Pb-Pf*CSP sporozoites (N=3). For panels E to G, top and bottom panels show Alexa Fluor 647 and Alexa Fluor 568 fluorescence, respectively. Statistical analysis comparing groups in (A) using unpaired t-test, in (E-F) using 2-way ANOVA with Tukey’s test for multiple comparisons, and in (G) using 2-way ANOVA with Dunnett’s test for multiple comparisons. Statistical significance indicated as *=P<0.05, **=P<0.01, ***=P<0.001, and ****=P<0.0001.

### The specificity of repeat-binding mAbs determines ctCSP accessibility on the sporozoite surface

Next, we measured the effect of repeat mAb epitope fine specificity on ctCSP unmasking and accessibility, quantified by binding of the previously characterized 1512 IgG^44^. We chose the well-characterized canonical mAbs binding the junctional (CIS43)^53^, minor (L9)^54^, and major repeats (311 and 317)^32^ of *Pf*CSP. Antibody binding to freshly isolated tg-*PbPf*CSP sporozoites was determined by flow cytometry following incubation with 10 µg/ml of Alexa Fluor568-labelled 1512 IgG and a serial dilution of Alexa Fluor647-labelled repeat-binding IgG (Fig. 3D). mAbs 311, 317 and L9 showed comparable concentration-dependent binding to sporozoites (Fig. 3E, top). The lower binding observed for CIS43 IgG could be attributed to CSP shedding, identifiable by the morphological changes of the sporozoite population (Fig. S5C)^55^. Across all concentrations, incubation with 311 or 317 IgG resulted in comparable ctCSP unmasking and 1512 IgG binding (Fig. 3E, bottom). Unmasking of ctCSP by mAbs 311 and 317 and subsequent 1512 IgG binding was highest at 0.3 µg/ml, and decreased at higher 311 and 317 IgG concentrations, which could potentially be due to antibody crowding on the sporozoite surface reducing accessibility to ctCSP (Fig. 3D, E). Unexpectedly, incubation with L9 IgG resulted in significantly higher 1512 IgG binding at 0.3-30 µg/ml (P<0.0001). Therefore, despite equivalent binding MFI to sporozoites (Fig. 3E, top), epitope fine-specificity of repeat-binding IgG can impact ctCSP unmasking and accessibility to mAb 1512 binding (Fig. 3E, bottom).

Diverse modes of antibody recognition of the CSP repeat region are also produced by Fab-Fab homotypic interactions often observed in V_H_3-33 mAbs and have been shown to contribute to protection in mice^31,51^. To determine the effect of homotypic Fab-Fab interactions on ctCSP accessibility, sporozoites were incubated with 1512 IgG-Alexa Fluor 568 and a serial dilution of three V_H_3-33 IgG (311, 356, and 239) and their homotypic-knockout mutants (311R, 356R, and 239R) labelled with Alexa Fluor 647 (Fig. 3F)^31^. At a low concentration, 0.3 µg/ml, V_H_3-33 IgG and their homotypic-knockouts bound sporozoites equivalently (Fig. 3F, top), but binding of 1512 IgG to sporozoites was significantly higher when incubated with WT V_H_3-33 IgG relative to their homotypic-knockouts (P<0.0001 for 311, P=0.0017 for 356, P=0.0081 for 239) (Fig. 3F, bottom). At a high concentration of 30 µg/ml, homotypic-knockouts showed higher binding to sporozoites relative to their WT V_H_3-33 IgGs (P<0.0001 for 356 and 239) (Fig. 3F top) and mAb 1512 binding was higher on sporozoites incubated with homotypic knockout IgGs (P<0.0001 for all pairs). These results suggest that homotypic Fab-Fab interactions are beneficial for ctCSP unmasking at lower mAb concentrations (Fig. 3F, bottom) but are unfavorable at high concentrations. At the higher IgG concentrations, homotypic Fab-Fab interactions may promote IgG packing on the sporozoite surface that reduces 1512 IgG binding by decreasing ctCSP unmasking or accessibility.

Lastly, we incubated sporozoites with 10 µg/ml 1512 IgG-Alexa Fluor568 and 3 µg/ml repeat-binding IgG-Alexa Fluor 647 across a 2-hour period (Fig. 3G). For all repeat-binding mAbs, nearly 50% of mAb binding took place within the sample processing time (t=0 min) and plateaued starting at t=15 min (Fig. 3G, top). Despite an initial temporal delay, observed in the minimal 1512 IgG binding detected at t=0, ctCSP unmasking by mAbs 311 and 317 takes place by t=15 min and plateaus thereafter (Fig. 3G, bottom). Binding of 1512 IgG continues to increase for sporozoites incubated with L9, surpassing 311 by t=30 and with essentially no binding in the presence of CIS43 throughout. Therefore, differential ctCSP unmasking by repeat-binding mAbs, affected by epitope fine-specificity and homotypic Fab-Fab interactions, occurs on a physiologically relevant timescale^56^.

### ctCSP mAbs induced by RTS,S vaccination are weakly protective from transgenic sporozoite challenge in mice

Next, we determined the efficacy of our extended MAL071 ctCSP mAb panel in reducing *Pb-Pf*CSP chimeric *P. berghei* parasite liver-burden *in vivo* (Fig. S6)^44^. In five independent experiments, positive control mAb 311 provided consistently high reduction in liver burden (95–98%), whereas anti-Pfs25 negative control mAb 1245 did not provide significant protection (Fig. 4). Displaying both inter- and intra-experimental variability, ctCSP mAbs performed inconsistently between experiments providing 5–43% reduction in liver burden, with individual mAbs significantly reducing parasite liver-burden in some experiments. To increase the statistical power of our analysis, normalized liver burden reduction was pooled from all five experiments (Fig. 4 bottom right, Fig. S6). Across all five experiments, all ctCSP mAbs produce a significant reduction in liver burden, albeit only weakly protective (23–38%) compared to mAb 311 (97%). Overall, mAbs targeting the ⍺-epitope or β-epitope regions protect equivalently from challenge with *P. berghei* sporozoites expressing matched-strain 3D7 *Pf*CSP. However, due to the strain-specificity of mAbs targeting the ⍺-epitope region, only mAbs against the β-epitope regions are expected to protect against non-3D7 strains.

**Figure 4.**
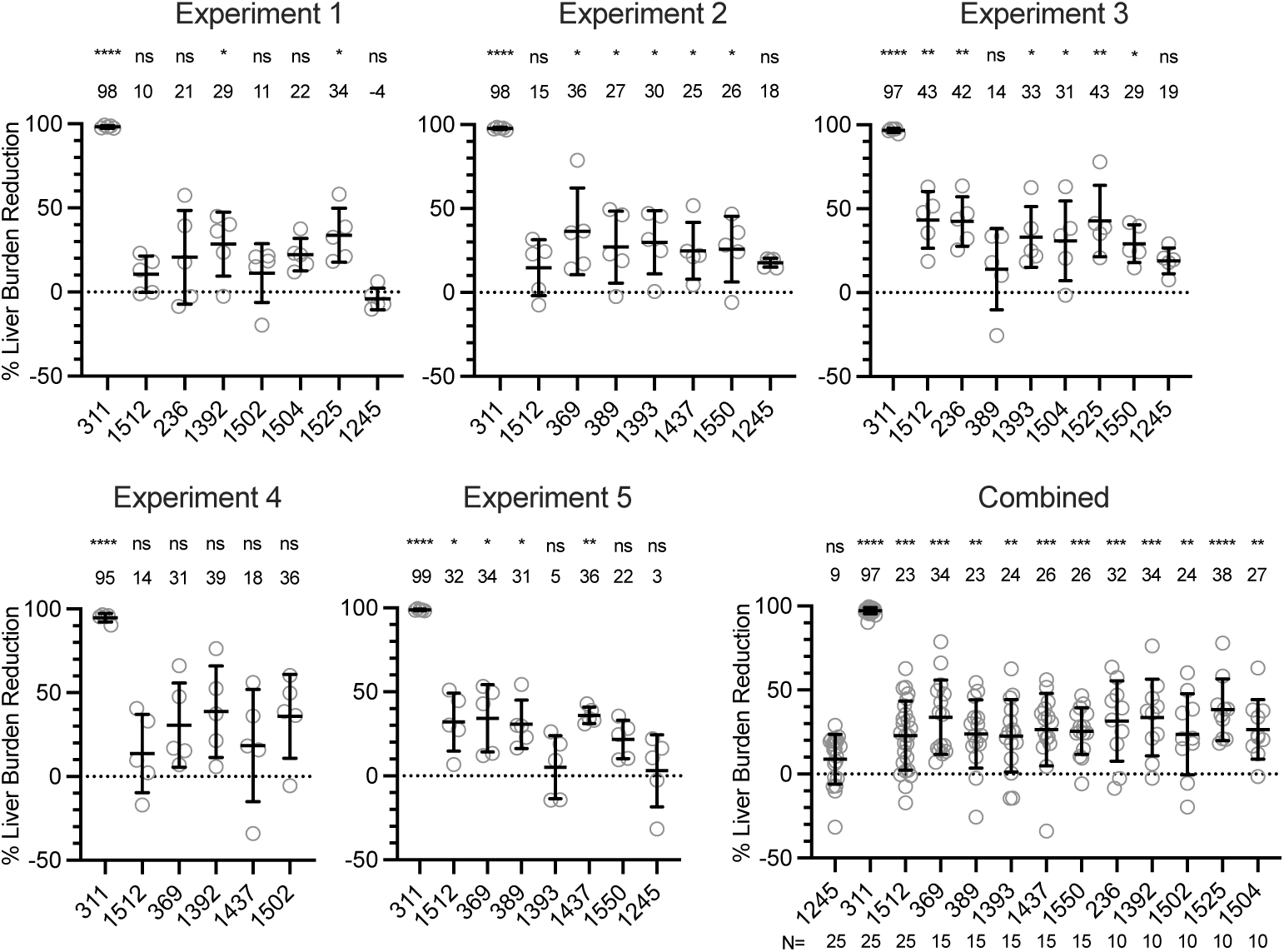
MAL071 mAbs are weakly protective from *in vivo* sporozoite challenge. Female C57Bl/6 mice were injected intravenously with IgG1 (300 µg/mouse) 16 hours prior to sporozoite challenge with 2000 transgenic *P. berghei* sporozoites expressing *Pf*CSP and luciferase. Forty-two hours post-challenge, mice were injected with D-luciferin and the liver sporozoite burden was measured as flux and normalized to the naïve control group. Percent liver burden reduction is shown above each group with pair-wise statistical analysis comparing groups against naïve using the Kruskal-Wallis non-parametric test (statistical significance indicated as *=P<0.05, **=P<0.01, ***=P<0.001, and ****=P<0.0001).

### mAb binding to cell surface ctCSP promote binding and downstream signaling through Fc-receptors

In addition to neutralizing their target through Fab binding to antigen, Fc-mediated effector functions are important mechanisms for antibody-mediated protection against many viral, bacterial, and parasitic pathogens^57^. The Fc-region of antibodies bound to *Pf*CSP can interact with Fc-receptors (FcR) on the surface of monocytes, neutrophils, and NK cells to mediate Fc-effector functions including phagocytosis and cellular cytotoxicity, with a key signaling role for Fc gamma receptor IIIa (FcɣRIIIa)^58–60^.

To dissect the effects of mAb epitope on Fc-mediated signaling, we used a diverse panel of 26 structurally and functionally characterized mAbs isolated from MAL071 vaccinees^31,33,34,61^ and patients immunized with irradiated sporozoites^53,54,62^ (Table S1). As a target, short CSP (sCSP) containing 19 NANP repeats was stably expressed on the surface of HEK293T cells (Fig. S7A). In addition to antibodies with a wild-type (WT) IgG1 backbone, we expressed Fc-variants with reduced binding to Fc receptors and complement C1q (L234A/L235A/P329G, LALAPG)^63^. Both WT and LALAPG antibodies bound cell-surface sCSP confirming that all target CSP epitopes are exposed in this format and Fc mutations do not diminish antigen binding (Fig. S7B). To determine FcɣRIIIa signaling of bound antibody, target cells expressing cell-surface sCSP were co-incubated with FcɣRIIIa-expressing Jurkat reporter cells. Only mAbs with LALAPG mutations in their Fc-region did not produce signal, confirming that mAb bound to target cells productively engage FcɣRIIIa specifically through their Fc region (Fig. S7C).

In our *in vitro* reporter assay, ctCSP mAbs bind cell-surface sCSP with a higher affinity and a significantly lower EC_50_ than repeat mAbs (p=0.0153), but signal through FcɣRIIIa with an equivalent EC_50_ (p=0.6869) (Fig. 5A). Nevertheless, mAbs to both epitopes demonstrate similar concentration-dependent binding and signaling dynamics, shown by the strong positive correlation in EC_50_ values (Pearson’s r=0.727, R^2^=0.528, p<0.0001) (Fig. 5A, right). As expected, mAbs to the repeat region showed significantly higher binding than those towards ctCSP (p<0.0001) (Fig. 5B). However, repeat-binding mAbs were more limited in their maximum FcɣRIIIa signal, ranging from 561–897 relative light units (RLU) with a mean of 722 RLU, while ctCSP mAbs showed a broader range of 488–1820 RLU with a mean of 1000 RLU (Fig. 5B). Overall, maximal signal was higher for ctCSP mAbs compared to repeat-binding mAbs (p=0.058), and there was no correlation between maximal binding and maximal FcɣRIIIa signal (Pearson’s r=-0.327, R^2^=0.107, p=0.103). Despite large intra-epitope variation in maximal FcɣRIIIa signal, the higher overall signal for ctCSP mAbs compared to repeat mAbs suggests that for *Pf*CSP overall FcɣRIIIa signaling is more dependent on epitope specificity than binding stoichiometry.

**Figure 5.**
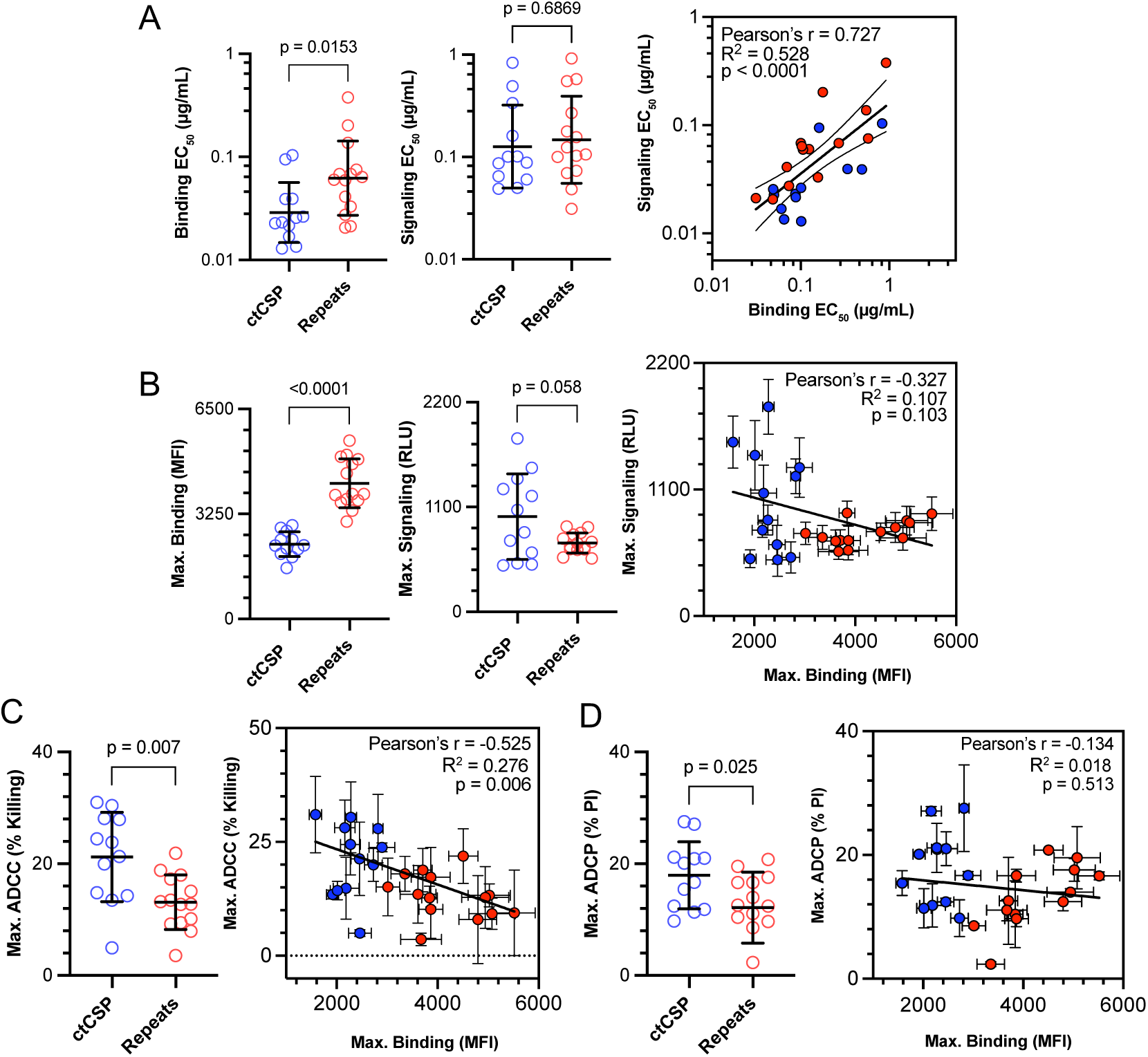
mAbs against ctCSP promote FcɣRIIIa signaling and mediate effector functions in primary cells. Target cells expressing membrane-bound sCSP (see Methods) were incubated with antibody titrations. Antibody binding was assessed by flow cytometry. FcɣRIIIa signaling was measured following co-culture of target cells with Jurkat reporter cells expressing luciferase under the FcɣRIIIa-controlled NFAT promoter, and FcɣRIIIa signal measured using luciferase substrate. A) EC50 and B) Maximum values for antibody binding and FcɣRIIIa signaling and correlation on the right panel (in all panels). To measure ADCC, target cells were co-cultured with primary Natural Killer (NK) cells isolated from human PBMCs (N=2 donors) and killing was measured using flow cytometry. To measure ADCP, recombinant sCSP-coated beads were co-incubated with primary neutrophils isolated from human PBMCs (N=2 donors) and bead uptake was measured using flow cytometry. C) Maximum ADCC (% killing) and D) Maximum ADCP (% bead uptake), and correlation on the right panel. All values were calculated using a 3-point logistic curve. Repeat mAbs are shown in red and ctCSP mAbs in blue. All values from N=4 (binding and FcɣRIIIa signaling) or N=2 (ADCC and ADCP, per donor) experimental replicates with standard deviation shown as error bars. Significance is measured in (A-D) with Welch’s t-test.

We next measured CSP-specific antibody-dependent cellular cytotoxicity (ADCC) by primary NK cells. Target cells expressing cell-surface sCSP were co-incubated with primary NK cells and loss of intracellular green-fluorescent protein (GFP) fluorescence (indicating productive NK cell degranulation) was quantified by flow cytometry. mAbs to ctCSP demonstrated a ∼2-fold higher ADCC compared to repeat-binding mAbs (p=0.007). Across all mAbs targeting both ctCSP and the repeat epitopes, increased binding to cell-surface sCSP was associated with a decrease in ADCC (Pearson’s r=-0.525, R^2^=0.279, p=0.006) (Fig. 5C). To determine neutrophil antibody-dependent cellular-phagocytosis (ADCP), primary neutrophils were co-incubated with recombinant sCSP-coated beads in the presence of IgG, and bead uptake was quantified by flow cytometry. In this assay, ctCSP mAbs promoted marginally higher ADCP than repeat-binding mAbs (p=0.025), but with no significant correlation between ADCP and maximal binding values (Pearson’s r=-0.134, R^2^=0.018, p=0.513) (Fig. 5D).

### Combining mAbs against multiple epitopes augments FcɣRIIIa signaling and ADCC

Within the context of RTS,S vaccination and next-generation immunogen design, ctCSP mAbs must be considered in conjunction with repeat-binding mAbs. Therefore, we determined the effect of combining mAbs against both ctCSP and repeat epitopes on FcɣRIIIa signaling and ADCC following incubation of target cells with an equivalent and saturating concentration, 3 µg/ml each of ctCSP and repeat mAbs (Fig. S8A and S8B). Normalized to the signal produced by ctCSP mAb alone, addition of repeat mAbs increased FcɣRIIIa signal for mAbs with weak signaling potency (e.g. mAb 236) and decreased signal for potent FcɣRIIIa-signaling mAbs (e.g. mAb 1393) (Fig. 6A). Normalized to the FcɣRIIIa signal of repeat-binding mAbs alone, addition of ctCSP mAb increased FcɣRIIIa signal in most combinations (Fig. 6A). Increase in FcɣRIIIa signal in combination with ctCSP mAbs was highest for mAbs 317 and L9. In particular, mAb 369 combined with L9 or 317 produced a 70% increase in FcɣRIIIa signal compared to repeat mAb alone. Using the definition of Excess over Bliss (i.e., a mathematical metric for the difference between the expected and actually observed signal) mAb combinations that include a ctCSP mAb with strong FcɣRIIIa signaling are not additive (1393, 1502, 1525), whereas combining a ctCSP mAb with repeat mAb of a similar maximal FcɣRIIIa signal (236, 369, 1512) produces an additive or weakly synergistic FcɣRIIIa signal (Fig. 6B).

**Figure 6.**
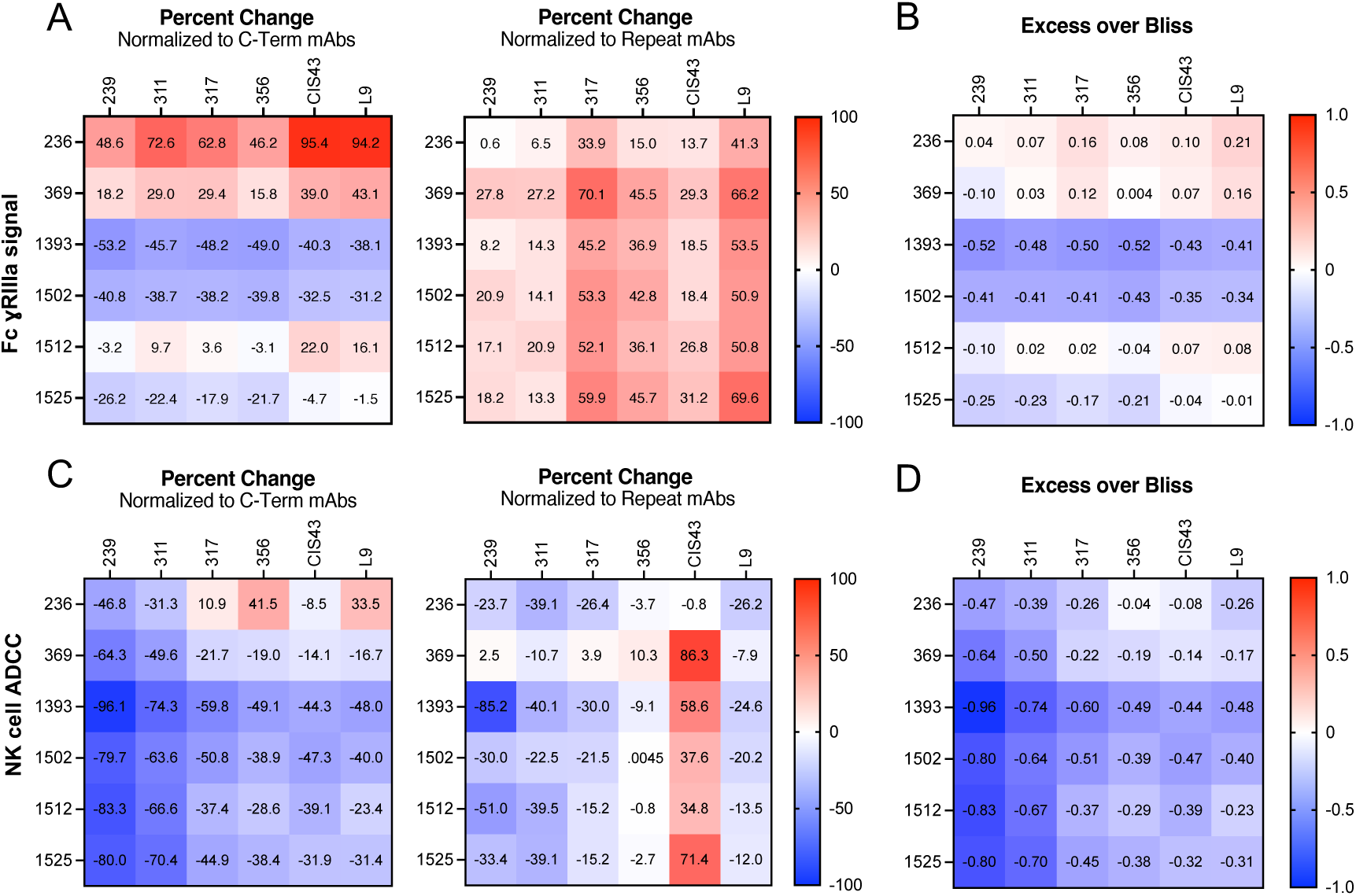
Combining mAbs against multiple CSP epitopes augments FcɣRIIIa signaling and alters ADCC. A) Percent change of FcɣRIIIa signal in Jurkat-Lucia NFAT-CD16 by combinations of mAbs at 3 µg/ml normalized to baseline of central repeat or ctCSP-binding mAb alone and B) Synergistic, additive, or antagonistic effects from mAb combination shown as Excess over Bliss using Bliss independence model. C) Percent change and D) Excess over Bliss of ADCC in primary NK cells by combinations of mAbs at 3 µg/ml. Positive values are shown in red and negative values are in blue.

In our ADCC assay using primary NK cells, most mAb combinations resulted in dampening of cellular cytotoxicity relative to both mAbs binding individually. Showing a similar trend to FcɣRIIIa signaling in NFAT-reporter Jurkat cells, addition of mAbs against the repeat region dampened the signal of ctCSP mAbs with potent ADCC (mAbs 1393, 1502, and 1525), and improved ADCC for mAb 236 with weak ADCC (Fig. 6C). The only repeat-binding mAb with an increase in ADCC upon addition of ctCSP-binding mAbs was CIS43, with mAbs 369 and 1525 producing an 86% and 71% increase in ADCC, respectively, over the baseline ADCC of mAb CIS43 alone (Fig. 6C). For each pair of mAbs, the resulting overall ADCC is lower than that of the most potent mAb in each combination, resulting in negative Excess over Bliss values, indicating near additivity (e.g. mAbs 369 and CIS43) or strong antagonism (e.g. mAbs 1393 and 239) (Fig. 6D). Therefore, depending on mAb fine specificity, ADCC of repeat-binding mAb may be enhanced by addition of ctCSP mAbs.

## Discussion

Since the discovery of the gene for CSP^8,11^, the 40-year-long journey of RTS,S/AS01_E_ development ^12,13^, and development of the similar but modified formulation of R21/Matrix-M^19,20^, the efficacy and potency of CSP-based vaccination has been attributed and associated with the antibody response to the central repeat epitope of CSP^22,27^. Together with the more recently identified junctional^53^ and minor repeat epitopes^54^, which are not included in RTS,S and R21, a myriad of structural, biophysical, and in vivo characterizations of mAbs have demonstrated the mechanisms for binding and potency of mAbs targeting the CSP repeat region^34,62,64,65^. In contrast, the humoral response to ctCSP has garnered less attention^42–45^, with no clear mechanism of action for the vaccine-induced antibody response to this domain. Here, crystal structures of 11 Fabs to ctCSP from the RTS,S/AS01_E_ MAL071 trial^33,36^ expanded the antigenic landscape to ctCSP and revealed multiple modes of strain-transcending binding to the conserved β-epitope region of ctCSP (Fig. 1C). In agreement with previous studies, we showed that mAb binding to ctCSP on live sporozoites is largely dependent on domain unmasking by some repeat antibodies, and further revealed the effects of antibody concentration, binding mode, and fine epitope specificity on ctCSP accessibility.

The antibody response to ctCSP is dominated by V_H_3-21 (and V_H_3-48) antibodies that bind the highly polymorphic ⍺-epitope of ctCSP^33,42,44,45^. In this study, we show that V_H_3-21, V_H_3-11, and V_H_4-50 mAbs engage the ⍺-epitope region through similar strain-specific interactions that are conserved in these different antibodies (Fig. 2A). RTS,S/AS01 trials that assess protection from CHMI using strain-matched parasites predict a higher vaccine efficacy than field trials on children and infants exposed to natural infection^16,23,36^. In one study, only 5.2% of infecting *Pf* strains matched the ⍺TSR sequence of the 3D7 vaccine strain^66^. Accounting for this strain mismatch improved RTS,S/AS01_E_ vaccine efficacy from 33% to 50% (p=0.04)^25^, specifically implicating responses to ctCSP polymorphic elements in vaccine-mediated protection.

The structural basis for strain-transcending ctCSP recognition, independently associated with vaccine-mediated protection, was first demonstrated by mAb 1512 binding to the ctCSP β-epitope region, which is conserved across ctCSP haplotypes^39,44^. However, mAb 1512 bound its β-epitope using an uncharacteristically long 21 amino acid HCDR3, compared to the average 14‒15 amino acid HCDR3 in the human repertoire. Broadly neutralizing antibodies (bnAbs) towards pathogens such as HIV often rely on long HCDR3s, with low precursor frequency in the human B-cell repertoire^67–69^. As a result, complex germline-targeting and multi-immunogen vaccination regimens have been devised to increase B-cell precursor frequencies and produce a robust bnAb response^68^. Our expanded panel of MAL071 mAb structures demonstrates a breadth of strain-transcending mAbs bound to ctCSP with shorter HCDR3s that are represented in the human antibody repertoire at higher prevalence (Table S1). Highlighting the plasticity of the antibody repertoire in binding this epitope, V_H_3-23 mAbs 1393, 1437, and 1550 with varied HCDR3 lengths pair with unique light chains to bind the β-epitope in distinct conformations (Fig. 2B). The extensive polar HCDR3 interactions of mAbs that bind to the β-epitope region contrasts to the HCDR1 and HCDR2-dominant polar interactions of V_H_3-21 mAbs to the ⍺-epitope region. Whereas the V_H_3-21 response to CSP is predominantly strain-specific, the full extent of strain-transcending responses to ctCSP requires antibody characterization beyond V-gene usage and consideration of HCDR3 D/J genes paired with determination of breadth and structure.

The coordinated exposure of ctCSP upon contact with hepatocytes during the phenotypic switch from migration to invasion provides a short timeframe for antibody binding and downstream effector functions^47^. This “just-in-time” exposure has been observed for other *Plasmodium* antigens such as RH5 *(P. falciparum* merozoites)^70^ and DBP (*P. vivax* merozoites)^71^ and could, in part, explain the weak inhibition observed for ctCSP mAbs on their own (Fig. 4). Antibody binding to the CSP central repeats extends this window of opportunity by prematurely unmasking ctCSP^43^. In this study, we have shown that premature unmasking of ctCSP by repeat-binding antibodies depends on antibody concentration, epitope fine-specificity, and Fab-Fab homotypic interactions^31^ (Fig. 3E–G). Therefore, the potential contribution of ctCSP responses to protection are not only determined by the quality and magnitude of the humoral response to ctCSP but, importantly, as a function of ctCSP unmasking by the antibody response to the central repeat region. As a corollary, rapid waning of antibody titers following vaccination and subsequent changes to ctCSP unmasking could produce temporal modulation of protection by antibodies targeting the individual CSP epitopes. In children vaccinated with RTS,S/AS01_E_, antibody responses to the NANP repeats wane more rapidly than ctCSP responses^41^. The protective association with ctCSP responses was strongest 18 months post-immunization and remained significant after adjusting for NANP responses^41^.

The increased association of ctCSP responses with protection after decay of NANP responses aligns with our results showing the highest ctCSP exposure at intermediate concentrations of some repeat-mAbs relative to the highest concentrations tested (Fig. 3E–F). Together, these findings may highlight a key role for ctCSP-mediated protection during the maintenance phase of the vaccine response, when B-cell responses to the central repeat region decline below a protective threshold.

Recent studies have highlighted Fc-dependent effector functions in protection from sporozoite invasion in *Pf*CSP repeat-binding mAbs^60,72^, and several serological analyses specifically highlight ctCSP-mediated effector functions as a correlate of protection^28,30,38,40,58^. Using a diverse mAb toolkit, we demonstrate that ctCSP mAbs signal through FcɣRIIIa more potently than repeat-binding mAbs in our *in vitro* assays (Fig. 5B). In primary human NK cell ADCC and neutrophil ADCP, ctCSP mAbs outperform repeat-binding mAbs (Fig. 5C and 5D), providing the first demonstration of a function for ctCSP mAbs that is in line with previous correlates of protection, and supports an Fc-dependent role for ctCSP mAbs in RTS,S/AS01_E_-mediated immunity.

Two factors that have been shown to negatively impact Fc-effector functions are an increase in epitope distance from the plasma membrane, particularly above ∼10 nm, and low antibody binding stoichiometry^73–75^. Including the ⍺TSR domain, disordered C-terminal linker, and the extended rod-like repeat region [∼20 nm in length for full-length CSP^76^ and ∼10 nm for sCSP in complex with V_H_3-33 Fab^31^ (Fig. S9)], binding of IgG to repeat epitopes may place Fc-regions at a sub-optimal distance from the membrane. In contrast with previous studies that used recombinant *Pf*CSP non-specifically cross-linked to ELISA plate, bead, or cell surface, we use recombinant and cell-surface sCSP that is specifically anchored at the C-terminus, which facilitates the resolution of epitope-specific antibody function by ensuring a biologically relevant and more uniform orientation of *Pf*CSP. The functional differences between the ctCSP and repeat epitopes would suggest that epitope distance from the membrane plays a dominant role over binding stoichiometry in modulating Fc-effector functions mediated by antibodies toward *Pf*CSP. However, our hypothesis does not rule out potential effects of differential Fc-region spacing or antibody crowding in our observations. Together, our *in vitro* assays also offer an attractive hypothesis into a potential evolutionary advantage of the immunodominant repeat region, whereby the high-stoichiometric binding of mAbs to the CSP repeat region provides a spatial buffer to distance Fc-receptors and soluble effectors from the parasite membrane.

Whereas mAbs in this study were chosen due to their well-characterized preference to individual epitopes within the *Pf*CSP repeat region, B-cell responses to *Pf*CSP repeat sequences are cross-reactive. Henceforth, investigation of ctCSP-mediated protection within the context of *Pf*CSP-based immunogens should prioritise the use of polyclonal sera to simulate a breadth of antibody affinities and binding modes that is produced by diverse epitopes, fine-specificities, and homotypic Fab-Fab interactions. The differential contribution of repeat-binding mAbs to ctCSP unmasking, Fc-signaling, and ADCC when combined with ctCSP mAbs supports the optimization of second-generation immunogens that include ctCSP to enhance and diversify Fc-mediated effector functions contributing to vaccine-mediated protection^30,58,77^. Furthermore, the superior unmasking of ctCSP by L9 (Fig. 3E) and improvement of CIS43 ADCC in combination with ctCSP mAb (Fig. 6C) encourage the development of next-generation immunogens that include ctCSP in combination with minor repeat and junctional sequences, which are both absent from RTS,S/AS01_E_ and R21/Matrix-M.

## Acknowledgements

We thank Fidel Zavala and Yevel Flores-Garcia (Johns Hopkins University) for conducting sporozoite challenges in mice to assess in vivo mAb liver-burden reduction. We thank the Insectary at UCSD and the Insectary and Parasitology Core Facilities at the Johns Hopkins Malaria Institute for providing *Pb-Pf*CSP and *Pb-Pf*ctCSP-infected mosquitos for flow cytometry and *in vivo* experiments. We thank the staff of the National Synchrotron Light Source II (NSLS-II) beamline 17-ID-2 (FMX), Stanford Synchrotron Radiation Lightsource (SSRL) beamlines BL12-1 and BL12-2, Advanced Photon Source beamlines 23-ID-B and 23-ID-D, and Argonne National Laboratory Advanced Light Source (ALS) beamlines 5.0.1 and 8.2.2 for assistance. This research used beamlines BL12-1 and BL12-2 of the Stanford Synchrotron Radiation Lightsource, SLAC National Accelerator Laboratory, which is supported by the U.S. Department of Energy, Office of Science, Office of Basic Energy Sciences under Contract No. DE-AC02-76SF00515. The SSRL Structural Molecular Biology Program is supported by the DOE Office of Biological and Environmental Research, and by the National Institutes of Health, National Institute of General Medical Sciences (P30GM133894). This research used beamline 17-ID-2 (FMX) of the National Synchrotron Light Source II, a US Department of Energy (DOE) Office of Science User Facility operated for the DOE Office of Science by Brookhaven National Laboratory under contract no. DE-SC0012704. The Center for BioMolecular Structure (CBMS) is primarily supported by the National Institutes of Health, National Institute of General Medical Sciences (NIGMS) through a Center Core P30 Grant (P30GM133893), and by the DOE Office of Biological and Environmental Research (KP1605010). This research was performed on APS beamlines 23-ID-B and 23-ID-D. GM/CA@APS has been funded by the National Cancer Institute (ACB-12002) and the National Institute of General Medical Sciences (AGM-12006, P30GM138396). The Eiger 16M detector was funded by NIH grant S10OD012289. This research used resources of the Advanced Photon Source, a U.S. Department of Energy (DOE) Office of Science User Facility operated for the DOE Office of Science by Argonne National Laboratory under Contract No. DE-AC02-06CH11357. This work was done on beamlines 5.0.1 and 8.2.2 of the Advanced Light Source, a DOE Office of Science User Facility under Contract No. DE-AC02-05CH11231, is supported in part by the ALS-ENABLE program funded by the National Institutes of Health, National Institute of General Medical Sciences, grant P30 GM124169-01.

This work was supported in part by the Gates Foundation INV-004923 and INV-056202 (I.A.W., D.R.B, T.F.R.). The conclusions and opinions expressed in this work are those of the authors alone and shall not be attributed to the Foundation. Under the grant conditions of the Foundation, a Creative Commons Attribution 4.0 License has already been assigned to the Author Accepted Manuscript version that might arise from this submission. Please note works submitted as a preprint have not undergone a peer review process. This work was also supported by PATH’s Center for Vaccine Innovation and Access, U.S. Agency for International Development (USAID) Innovations in Malaria Vaccine Development Contract Number 7200AA20C00017. This work was partially supported by a Skaggs-Oxford Graduate Fellowship to R.M. The funders did not play any role in the study design, data collection and analysis, decision to publish, or preparation of the manuscript. The content of this publication is solely the responsibility of the authors and do not necessarily represent the official views of the Gates Foundation, PATH, USAID, or NIH.

## Author contributions

Conceptualization: R.M., I.B., L.H., and I.A.W. Formal analysis: R.M., I.B., N.B., R.L.S., X.Z. Antibody sequences: E.L., R.S.M., K.L.W., D.E.E., C.F.O. Funding acquisition: E.A.W., C.R.K., D.R.B., T.F.R., L.H., I.A.W. Investigation: R.M., I.B., N.B., G.G.-P, M.J., T.Z., M.V.B., J.N., K.G., J.Z., R.L.S., X.Z. Methodology: R.M., I.B., N.B., G.G.-P, M.V.B., L.H. Supervision: L.H, I.A.W., R.M. Writing – original draft: R.M., I.B., L.H., I.A.W. Writing – review & editing: all authors.

## Materials availability

All unique reagents generated in this study are available to academic researchers from Ian A. Wilson with a completed materials transfer agreement

## Data availability

Crystal structures of the Fab-⍺TSR complexes have been deposited in the Protein Data Bank with accession codes: 9O8O (Fab 367), 9NI2 (Fab 369), 9NI1 (Fab 389), 9NCY (Fab 1392), 9NIY (Fab 1393), 9NJ4 (Fab 1437), 9NHY (Fab 1502), 9O18 (Fab 1504), 9NDM (Fab 1525), 9NHV (Fab 1534), 9NIW (Fab 1550).

## Competing interests

The authors declare no competing interests.

## Supplementary material

Materials and methods, Figures S1-S9, Tables S1-S6, supplementary references

## Notes

### Competing Interest Statement

The authors have declared no competing interest.

