## Supplemental File for "Structure and functional diversity of antibodies targeting the *P. falciparum* circumsporozoite protein C-terminal domain"

### Materials and methods

#### Ethics statement

The assays using mice were performed in strict accordance with the recommendations in the Guide for the Care and Use of Laboratory Animals of the National Institutes of Health. The protocol was approved by the Animal Care and Use Committee of the Johns Hopkins University, protocol number MO18H419.

#### Sequence and structural analysis

MAL071 antibody sequences were originally isolated, characterized, and described by Williams et al.[33] and provided by PATH's Center for Vaccine Innovation and Access. Antibody sequences were analyzed with Ig-BLAST[87] and abYsis[88]. Kabat numbering is used throughout, with superscript H and L to distinguish antibody heavy and light chain residues from non-antibody residues. Buried surface area was calculated using the Molecular Surface package (MS)[89] using a 1.7 Å probe radius, protein-protein interactions were determined using the PDBePISA server of EMBL-EBL, and water-mediated interactions were determined manually in Coot[90]. Structural figures were produced using Mac PyMOL (Schrödinger LLC) Version 2.5.1 and graphical data were plotted using GraphPad Prism.

#### rsCSP and αTSR expression in *E. coli*

CSP αTSR domain (*P. falciparum* 3D7 amino acids 310-376) with a $His_6$-tag (for X-ray crystallography) or with a C-terminal avi-tag and $His_6$-tag (for epitope binning) were inserted into pET28a expression vector and expressed in *E. coli* SHUFFLE competent cells as outlined previously[44]. rsCSP with a C-terminal avi-tag and dual N- and C-terminal $His_6$-tags was inserted into pET28a expression vector and expressed in *E. coli* SHUFFLE competent cells as detailed previously[32,91]. For binding breadth analysis, 15 ctCSP alleles capturing various polymorphisms in Th2R and Th3R epitopes[44] with a C-terminal strep tag in a pET28a vector, were expressed in *E. coli* SHUFFLE cells. Cultures were induced at OD 0.6–0.8 with 1 mM isopropyl β-D-1-thiogalactopyranoside and grown at 18°C overnight. Bacterial cell pellets were ruptured using a EmultiFlex-C3 microfluidizer for 10 min on ice, and supernatants were clarified by centrifugation at 26,000 xg for 30 min at 4°C, loaded onto a StrepTrap HP column, washed with phosphate-buffered saline (PBS), and eluted with PBS +2.5 nM D-desthiobiotin. Relevant elution fractions were dialyzed in PBS overnight at 4°C, purified by size exclusion chromatography on a HiLoad™ Superdex75 16/90 column (GE Healthcare), aliquoted and stored at -80°C.

#### IgG and Fab expression in mammalian cells

Constructs for mammalian expression were subcloned into a pCMV3 vector with a kappa light chain leader sequence or a AbVec1 with an IL2 leader sequence. For Fab expression, a stop codon was introduced between the $C_H1$ and hinge region of the IgG1 sequence. For X-ray crystallography of Fab1534, some light chain residues of the $C_K$ domain were mutated to improve crystal packing and crystallization[92]. Antibody light and heavy chain MidiPrep and MaxiPrep DNA were purified separately (Takara Bio Inc.) following the manufacturer's protocols and transfected at a 1:1 vector mass ratio. For X-ray crystallographic analysis and protection studies in mice, Fab and IgG1 were expressed in ExpiCHO cells using Lipofectamine transfection reagent (Invitrogen) following the manufacturer's protocol. Cells were cultured in ExpiCHO expression medium (Gibco) and supernatant harvested after 10-12 days. For Fc-receptor binding studies, IgG were expressed in HEK293F cells cultured in Freestyle293 expression media (Gibco) or Expi293F cells cultured in Expi293™ Expression

Medium (Gibco). For HEK293F cell transfection, DNA and 3x v/v PEI-MAX 40,000 (Polysciences) individually diluted in OptiMEM (Gibco) were mixed and allowed to incubate at RT for 20–30 min before addition to cells. Expi293F cells were transfected with ExpiFectamine™ 293 Transfection Kit (Gibco) following the manufacturer's protocol. For both HEK293F and Expi293F cell cultures, supernatants were harvested after 6–7 days. Fabs were purified using a HiTrap Protein G HP column (GE Healthcare) and size exclusion chromatography on Superdex200 16/90 or Superdex75 16/60 columns (GE Healthcare) equilibrated with Tris-buffered saline (TBS, 50 mM Tris pH 8.0, 137 mM NaCl, 3.6 mM KCl). IgG were purified using a HiTrap Protein A HP column (GE Healthcare) and size exclusion chromatography on Superdex200 16/90 column (GE Healthcare) equilibrated with dPBS (Gibco).

***In vitro* Avi-tag protein biotinylation**

Avi-tagged αTSR domain and rsCSP purified as detailed above were dialyzed in 4°C overnight using SnakeSkin dialysis tubing (Thermo Fisher Scientific) in 200 mM KCl and 50 mM Tris pH 8 buffer. Protein samples were biotinylated for 1 hour at RT with addition of 500 µl Biomix B and 25 µl BirA (5 mg/ml stock) per 5 ml protein sample adjusted to 40 µM. Excess biotin was removed by size exclusion chromatography on a Superdex75 16/60 column or Superdex200 16/90 column (GE Healthcare). Elution fractions were pooled and stored at -80°C.

**Epitope binning by bio-layer interferometry**

Epitope binning was performed using bio-layer interferometry on an Octet HTX (FortéBio). Biotinylated αTSR domain was loaded onto streptavidin sensors (Sartorius 18-5019) at 60 nM in running buffer (PBS + 0.05% Tween 20). Loaded sensors were then dipped into running buffer for 60 seconds followed by saturating mAb at 30 µg/ml for 600 seconds. Sensors were then dipped in running buffer for 60 seconds followed by the second competing mAb at 10 µg/ml for 300 seconds. Epitope binning experiments were run bidirectionally with each mAb. Data were analyzed using FortéBio analysis software. Normalized response, in nm shift, of the saturating mAb was plotted against the normalized response of the competing mAb to generate the competition matrix.

**Surface Plasmon Resonance**

Affinity experiments were performed on a Biacore 8k at 25°C. All experiments were carried out with a flow rate of 30 µl/min in a mobile phase of HBS-EP+ (0.01 M HEPES pH 7.4, 0.15 M NaCl, 3 mM EDTA, 0.0005% v/v Surfactant P20). Anti-Human IgG (Fc) antibody was immobilized via standard NHS/EDC coupling to a Series S CM-5 sensor chip. Each mAb was injected over the chip followed by a wait period to normalize captured response units (RU). A concentration series of each ctCSP variant was injected across the antibody and control surface for 2 min, followed by a 5000–8000 second dissociation phase. Regeneration of the surface in between injections of ctCSP variants was achieved with a single, 120 s injection of 3 M $MgCl_2$. Kinetic analysis of each reference subtracted injection series was performed using the BIAEvaluation software (Cytiva). All sensorgrams were fit to a 1:1 (Langmuir) binding model of interaction.

**Structure determination by X-ray crystallography**

For crystallographic analysis, purified Fab was combined with a molar excess of αTSR domain. Fab-αTSR complexes were purified by size exclusion chromatography on a HiLoad™ Superdex200 or Superdex75 16/600 column (GE Healthcare) equilibrated with TBS and an excess of αTSR was confirmed by the presence of an elution peak corresponding to unbound αTSR. mAb 1502 did not express in Fab format and $F(ab')_2$ was produced by digestion of 1502 IgG with IdeS protease (produced in-house) at room temperature, purified by size exclusion chromatography on a HiLoad™ Superdex200 16/90 column, with $F(ab')_2$-αTSR complex produced as described above. Relevant elution fractions were pooled and concentrated with 3 kDa or 10 kDa cut-off centrifugal filters (Millipore Amicon or Sartorius Vivaspin) to concentrations of 10–30 mg/ml. In several samples, Protein G domain III from Streptococcus (produced in-house) was added in equimolar amounts as crystallization chaperone. Crystallization screens were conducted with our high-throughput robotic Rigaku CrystalMation system at The Scripps Research Institute by sitting drop vapor diffusion with all Fab-αTSR complexes producing crystals in multiple conditions. Crystals were looped in mother liquor supplemented with cryoprotectant and stored in liquid nitrogen until data collection. For the datasets shown, conditions that produced the best diffracting crystals are summarized in Table S5. Datasets were collected at APS 23-ID-B, APS 23-ID-D, ALS 501, ALS 8.2.2, SSRL 12-1, SSRL 12-2, and NSLS-II 17-ID-2 by rotating crystals for a full 360°, collecting diffraction data every 0.2° to produce 1800 frames per dataset. Diffraction data for Fab369-αTSR were processed by XDS[93]. All other diffraction data were processed using HKL2000[94]. Structures were determined through molecular replacement in Phaser[95] using variable domain search models produced by SAbPred[96], variable domain fragments lacking CDRs, and constant domain search models for kappa or lambda light chain Fabs. To ensure correct placement of the αTSR domain, a separate search model was used. Structures were refined in REFMAC5[97] and PHENIX[98], and built and manually refined in Coot[90]. For datasets containing multiple Fab-αTSR complexes in the asymmetric unit, space group assignment was confirmed with Zanuda[99]. Data collection and refinement statistics are summarized in Tables S2–4. In the Fab367-αTSR structure, limited crystal contacts made by the αTSR domain in the crystal lattice resulted in high disorder and poor regional electron density. These features are reflected by the high Wilson B and individual B values for the αTSR domain as compared to the Fab, which are listed separately for all datasets in Tables S2–4.

**Sporozoite culture and isolation**

Sporozoites used for flow cytometry and *in vivo* challenge, termed tg*Pb-Pf*CSP are transgenic *P. berghei-Pf*CSP sporozoites (strain ANKA 676m1cl1, MRA868 background) expressing full-length *Pf*CSP strain 3D7 and a green fluorescent protein/luciferase fusion protein. A sterile complete RPMI 1640 media (Gibco) containing 5,000–10,000 sporozoites was injected into mice via retro-orbital administration. Parasitemia was monitored daily from day 6 post-infection using Giesma staining solution (CAS 51811-82-6). Mice were anesthetized with a ketamine/xylazine solution at 5–10% parasitemia and 1–2% gametocytemia and fed to female *Anopheles stephensi* mosquitoes that were maintained at approximately 21°C and 75% relative humidity. For flow cytometric analyses and intravenous (IV) challenge, day 21 infected mosquitoes were immobilized at 4°C for 5 min and sequentially washed in 70% ethanol, PBS, and complete RPMI. Salivary glands were dissected using insulin syringes and kept on ice. Sporozoites were purified with a glass homogenizer and filtered through a 20 µm filter (EMD Millipore). Sporozoite solutions were then concentrated and allowed to settle for 10 min before being counted on a hemocytometer (iNCYTO C-Chip Neubauer Improved, DHC-N01).

**Flow cytometry of mAb binding to sporozoites**

Freshly isolated salivary gland sporozoites were stained with SYBR Green I (Invitrogen) at a final concentration of 1:2,000 for 30–60 min on ice. Due to variable yields of sporozoites from mosquito salivary gland dissection, approximately 8,000–30,000 sporozoites were used per sample for flow cytometric analysis. To remove salivary gland debris, sporozoites were thoroughly resuspended in DMEM (Dulbecco's Modified Eagle Medium, Corning 15013CV), applied onto a 17% w/v solution of Accudenz in deionized water, and centrifuged for 20 min at 4°C 2500 xg at 0 deceleration[100]. Accudenz interface containing sporozoites were aspirated and centrifuged for 5 min at 21,000 xg and the sporozoite pellet resuspended in an appropriate volume of dPBS + 1% BSA. mAbs were directly labelled with Alexa Fluor 568 (Thermo Fisher Scientific A20184) and Alexa Fluor 647 (Thermo Fisher Scientific A20186) following the manufacturer's protocol or used unlabeled and diluted as necessary in dPBS + 1% BSA. 50 µl sporozoites were added to 50 µl IgG dilutions and incubated at room temperature (RT). For experiments in figure 4B, sporozoites were stained for 60 min, and for experiments in figures 4C, E, and F, for 30–45 min to reduce CSP shedding. Sporozoites were washed with dPBS + 0.25% BSA and resuspended in PBS + 1% BSA for analysis on the BioRad ZE5 Cell Analyzer. Raw data were analyzed on FloJo (Version 10.9.0), and data were graphed and analyzed in GraphPad Prism (v9.4.1).

***In vivo* mouse challenge using transgenic *Pb-Pf*CSP-GFP/Luc sporozoites**
Female C57Bl/6 mice (6–8 weeks old) were injected intravenously with endotoxin-free monoclonal antibodies 16 hours prior to sporozoite challenge. For intravenous (IV) sporozoite challenge, mice were injected with 2,000 chimeric *P. berghei Pf*CSP or *P. berghei Pf*ctCSP sporozoites. Forty-two hours after challenge, mice were injected with 100 µl of D-luciferin (30 mg/ml), anesthetized with isoflurane and the liver imaged with the IVIS Spectrum to measure the bioluminescence expressed by the chimeric parasites. Results were provided as Total Flux (photons/sec) and percent liver burden reduction was calculated as 100*(1-(flux/geomean (Naïve))). For pairwise analysis of significance, the Kruskal-Wallis test was used to compare mean liver burden reduction to the naïve group. Individual P values are summarized * P<0.05, ** P<0.01, *** P<0.001), and **** P<0.0001.

**Generation of short CSP expressing HEK293T cell-line**
The open reading frame of sCSP (Met1–Ser375 of 3D7 CSP with a truncated repeat domain containing 19 NANP repeats) was subcloned into phCMV3 vector containing a C-terminal vesicular stomatitis virus glycoprotein G (VSVG) linker and transmembrane domain followed by self-cleaving green fluorescent protein (phCMV3-sCSP-VSVG-GFP) to validate expression. The sCSP-eGFP amplicon was then sub-cloned into a pLENTI vector for stable transfection. One day prior to transfection, 1.5 million HEK293T cells (ATCC CRL-3216) were seeded in a 10 cm culture dish in 10 ml of 293T transfection medium consisting of DMEM (Corning 15013CV) supplemented with 10% fetal bovine serum (FBS) and 2 mM L-glutamine (Corning 25005CI). HEK293T cells were transfected with 24 µg of maxiprep sCSP DNA (Qiagen 12963) using 42 µl of Lipofectamine 2000 (Invitrogen 11668027) following the manufacturer's protocol and cultured overnight. The next day, culture medium was replaced with 10 ml of 293T culture medium consisting of DMEM supplemented with 10% FBS, 2 mM L-glutamine, and 100 IU/ml-100 µg/ml Penicillin-Streptomycin (Corning 30002CI). The following day, cells were rinsed with PBS, detached using PBS supplemented with 2 mM EDTA for 3 min, and transferred to a new 10 cm culture dish in 10 ml of 293T selection medium consisting of DMEM supplemented with 10% FBS, 2 mM L-glutamine, 100 IU/ml-100 µg/ml Penicillin-Streptomycin, and 10 µg/ml Puromycin (Sigma P9620). After one week of culture under selection, positive cells were enriched twice by fluorescence-activated cell sorting (FACS, BD FACSMelody), gating on the top 1% cells staining for both GFP and mAb 311, with cellular expansion in selection media between enrichments. A monoclonal cell line was obtained by limiting dilution. Cells were detached using PBS supplemented with 2 mM EDTA and 10 mM HEPES and resuspended to a density of 2000 cells/ml in 293T culture medium. 200 µl of cell suspension were added to the top left well of a tissue culture-treated 96 well plate and serially diluted in 293T selection medium 1:2 down the column which was then

serially diluted 1:2 across the plate. After two weeks of culture, monoclonal GFP-positive colonies were selected for expansion.

**Antibody binding to sCSP-expressing HEK293T cells**

Antibodies were centrifuged for 15 min at 20,000 xg at 4°C, filtered using 0.22 µm filter tubes (Costar UX0193730) and diluted as required in IMDM (Iscove's Modified Dulbecco's Medium, Gibco 12440053) supplemented with 10% FBS, 2 mM L-glutamine, and 100 IU/ml-100 µg/ml Penicillin-Streptomycin (assay medium). Antibodies were diluted alone or in combination, and 50 µl of antibody dilution was transferred to tissue culture-treated 96-well plates (Corning 3595). sCSP expressing HEK293T target cells were detached using PBS supplemented with 2 mM EDTA and 10 mM HEPES and collected in assay medium. Target cells were centrifuged at 500 xg for 5 min, washed with signaling assay medium, centrifuged, and resuspended at a density of 2 million cells/ml in signaling assay medium. 50 µl of target cell suspension was transferred to antibody dilutions and incubated for 2 hours at 37°C with 5% $CO_2$ in 95% humidity. To quantify antibody binding, sCSP-expressing HEK293T target cells were washed 3 times with FACS Buffer (PBS supplemented with 2% FBS and 1mM EDTA). Allophycocyanin (APC)-labelled anti-Human IgG AffiniPure $F(ab')_2$ Fragment Donkey Anti-Human IgG (H+L; Jackson ImmunoResearch, 709-136-149) was diluted 1:200 in FACS buffer. 20 µl of the diluted Anti-Hu IgG was then added to each well for a final concentration of 5 µg/ml and incubated for 30 min at 4°C in the dark. After incubation, cells were washed 3 times with FACS Buffer and fixed for 15 min at 4°C using 50 µl of Cytofix/Cytoperm (BD 554722). After fixation, cells were washed 3 times with FACS Buffer. GFP and APC fluorescence were measured by flow cytometry on a BD ZE5 instrument with limits of 5,000 events or 60 seconds.

**NFAT-CD16 FcγRIIIa *in vitro* signaling assay**

For FcγRIIIa in vitro signaling, Jurkat-Lucia NFAT-CD16 (Invivogen jktl-nfat-cd16) effector cells were collected from suspension by centrifugation at 300 xg for 5 min, resuspended in assay medium, centrifuged, and resuspended at a density of 2 million cells/ml in signaling assay medium. 100 µl of effector cell suspension were transferred to assay plate containing target cells and antibodies prepared as detailed above. Assay plate was incubated overnight at 37°C with 5% $CO_2$ in 95% humidity. After overnight incubation, QUANTI-Luc™ 4 Reagent (25x) and QUANTI-Luc™ 4 Stabilizer (20x) (Invivogen rep-qlc4lg5) were prepared as per manufacturer's instructions. In a white half-well 96-well plate (Corning 3688), 50 µl of the assay plate supernatant were transferred to 50 µl of QUANTI-Luc™ 4 substrate and luminescence was immediately measured using BioTek Synergy H1 plate reader. For both assays, half-maximal effective concentrations were interpolated using a Hill-curve-based non-linear curve fitting model (GraphPad Prism 10) and maximal binding was set at 3 µg/ml.

**ADCC Natural Killer cell assays**

Antibodies were centrifuged at 20,000 xg for 15 min at 4°C, filtered using 0.22 µm filter tubes (Costar UX0193730), and diluted to 40 µg/ml in PBMC assay medium - RPMI1640 (Corning 15-040-CV) supplemented with 10% FBS, 2mM L-glutamine, and 100 IU/ml-100 µg/ml Penicillin-Streptomycin. In a dilution plate, each antibody was serially diluted across the plate using half-log dilutions. 50 µl of each antibody dilution were transferred to tissue culture-treated 96 well plates (Corning 3799). HEK293T-sCSP cells were detached in 1x PBS supplemented with 2mM EDTA and 10mM HEPES, collected in DMEM supplemented with 10% FBS, 2mM L- glutamine, and 100 IU/ml-100 µg/ml Penicillin-Streptomycin, washed with PBMC assay medium, and then resuspended at $2x10^6$ cells/ml in PBMC assay medium. 50 µl of target cell suspension was transferred to the assay plate containing antibody dilutions and incubated for 2 hours at 37°C with 5% $CO_2$ in 95% humidity. Natural Killer (NK) cells were isolated from frozen PBMCs using STEMCELL EasySep Human NK Cell Isolation Kit (STEMCELL 17955) following manufacturer's protocol. Isolated NKs were resuspended to a density of 2 million cells/ml in PBMC assay medium, and 100 µl of the isolated NK suspension was transferred to assay plate containing target cells and antibodies. Assay plate was incubated overnight at 37°C with 5% $CO_2$ in 95% humidity. The following day, cells were centrifuged at 500 xg for 5 min, supernatant was removed, cells were detached from wells and transferred to a non-TC treated plate (Corning 3788). Wells were washed with FACS buffer 3 times and fixed using 50 µl of Cytofix/Cytoperm (BD 554722) for 15 min at 4°C. After fixation, cells were washed 3 times with FACS buffer and resuspended in 150 µl of FACS buffer. GFP fluorescence was measured via flow cytometry (Bio-Rad ZE5) gated on target cells using FSC-488 Area/ SSC-488 Area, single cells using FSC-488 Area/ FSC-488 Height, and GFP+ using GFP-488 (525/35)/ FSC-488 Area. Acquisition limits were set for 5,000 events in the first gate or 40 µl of total volume. Percent killing was calculated as (100% · [1-(%GFP+ w/o mAb - %GFP+ w/mAb)/%GFP+ w/o mAb], where %GFP+ w/o mAb is the percent of GFP-positive target cells incubated without antibody and %GFP+ w/mAb is the percent of GFP-positive target cells that were incubated with a given antibody and given concentration.

**ADCP Neutrophil assay**

Per 96 well plate sample, 10 µl of 1 µm FITC-labeled FluoSpheres NeutrAvidin Microspheres (Invitrogen #F8776) were washed 3 times with 1 ml of PBS via centrifugation for 10 min at 5,000 xg and finally resuspended in 300 µl of PBS. rsCSP was added to a concentration of 30 µg/ml, gently agitated in the dark for 2 hours at 4°C, washed 3 times with PBS and resuspended in PBMC assay medium. Filtered and centrifuged antibodies were diluted to 40 µg/ml in PBMC assay medium. In a dilution plate, each antibody was serially diluted across the plate using half-log dilutions and 20µl of antibody dilutions were transferred to a 96 well plate (Corning 3788) together with 10µl of rsCSP-coupled beads and incubated for 2 hours at 37°C with 5% $CO_2$ in 95% humidity. During incubation, neutrophils were isolated using the EasySep™ Direct Human Neutrophil Isolation Kit (StemCell 19666) following the manufacturer's protocol. Neutrophils were washed once in PBMC assay medium by centrifugation for 5 min at 300 x*g* and resuspended at a density of $1x10^6$ cells/ml in PBMC assay medium. 50 µl of Neutrophil cell suspension were transferred to opsonized rsCSP-coated beads and cultured for 30 min at 37°C with 5% $CO_2$ in 95% humidity as previously described in Stefanutti *et al.* (2025) [60]. After incubation, cells were washed 3 times with FACS Buffer via centrifugation for 5 min at 500 x*g*. Cells were stained for CD16 (BioLegend 360730) for 30 min at 4°C in the dark. After staining, cells were washed 3 times with PBS then fixed with 50 µl of Cytofix/Cytoperm for 15 min at 4°C in the dark. After incubation, cells were washed twice with FACS Buffer and resuspended with 150 µl of FACS buffer. GFP fluorescence was measured via flow cytometry (Bio-Rad ZE5) gated on target cells using FSC-488 Area/ SSC-488 Area, single cells using FSC-488 Area/ FSC-488 Height, FSC-488 Area/ CD16-BV510-405 (525/50), and GFP+ using GFP-488 (525/35)/ FSC-488 Area. Acquisition limits were set for 5,000 events in the first gate or 40 µl of total volume. The percent Phagocytic Index (% PI) was calculated as 100% · [(Phagocytic Index of % $GFP^+$ w mAb - Phagocytic Index of % $GFP^+$ w/o mAb) / Phagocytic Index of % $GFP^+$ w/o mAb], where Phagocytic Index of % $GFP^+$ w/o mAb is the percent of GFP-positive cells multiplied by the MFI incubated without antibody and Phagocytic Index of % $GFP^+$ w/mAb is the percent of GFP-positive cells multiplied by the MFI that were incubated with a given antibody and given concentration.

### Supplementary figures

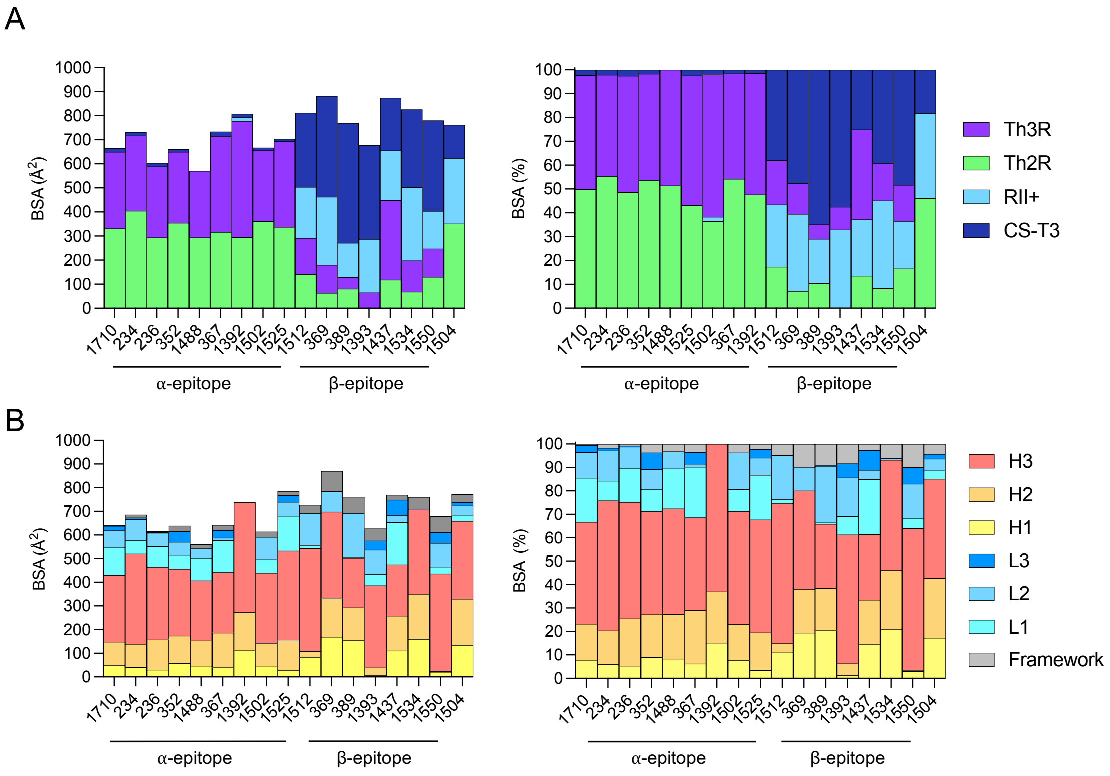


**Figure S1. Buried surface area of Fab-αTSR complexes.** (A) Buried surface area and percent formed in Fab-αTSR complexes on the αTSR surface grouped by epitope (Th2R 310–327, RII$^{+}$ 330–348, Th3R 348–363, and CS-T3 364–375). (B) Buried surface area (BSA) and percent BSA formed in Fab-αTSR complexes on the Fab surface grouped by framework and complementarity determining regions (H1–3 and L1–3 by Kabat numbering).

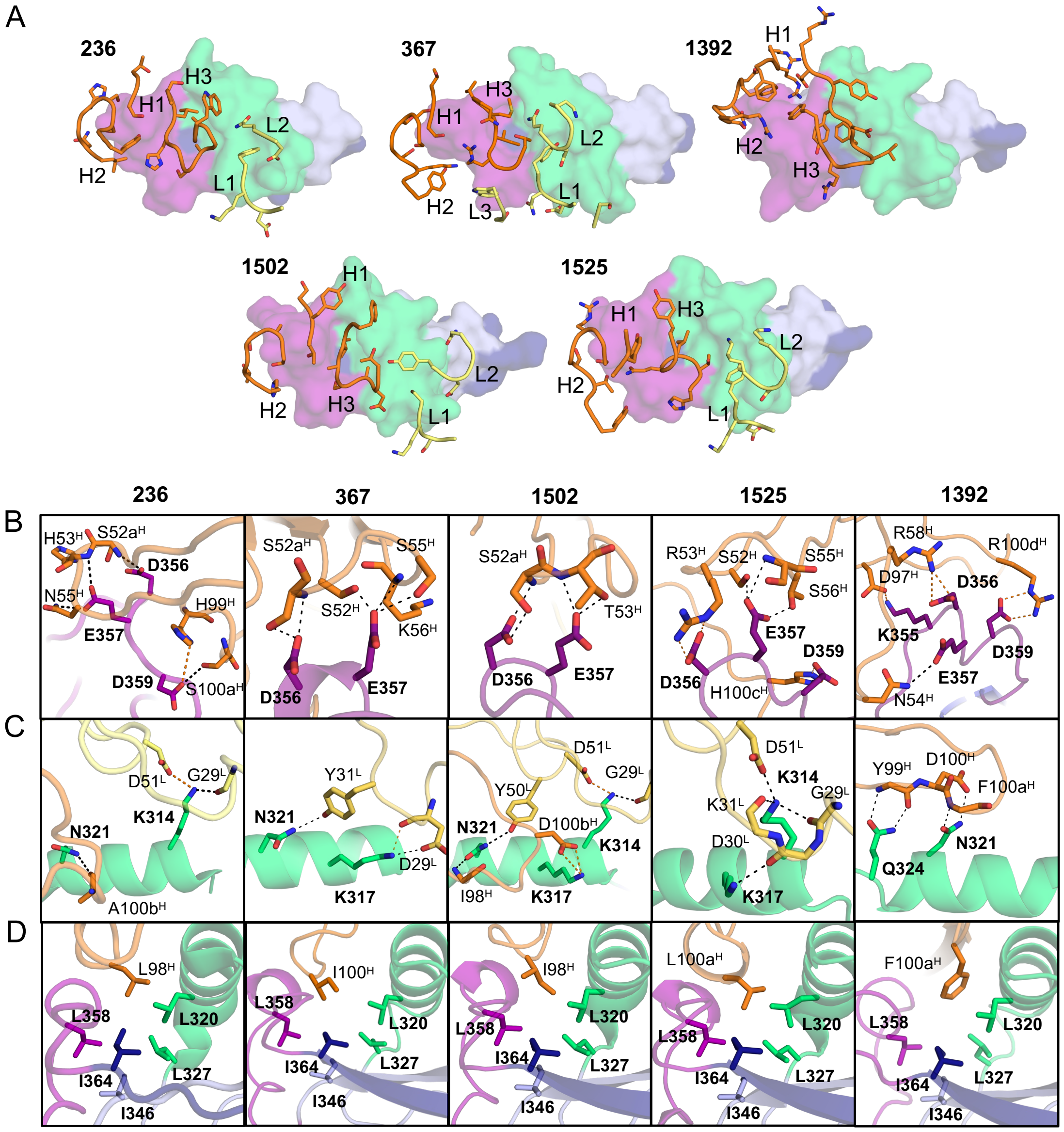


**Figure S2. Paratopes and molecular interactions of α-epitope binding Fabs with *Pf*CSP αTSR domain.** A) Surface representation of αTSR surface with paratopes defined as residues forming $>5Å^2$ BSA are shown as sticks, with CDRs labeled H1–3 and L1–3. Conserved interactions of α-epitope binding Fabs with B) highly polymorphic residues in the Th3R CS-flap and C) Th2R helix. D) Hydrophobic HCDR3 residues buried in the αTSR hydrophobic core. Fab heavy chain (orange) and light chain (yellow) backbone are shown in cartoon representation. The αTSR domain is shown with Th2R in green, Th3R in magenta, TSR-homology domain in blue, and CS-T3 in dark blue. Previously characterized Fab 236 (PDB:7RXJ) is shown for reference.

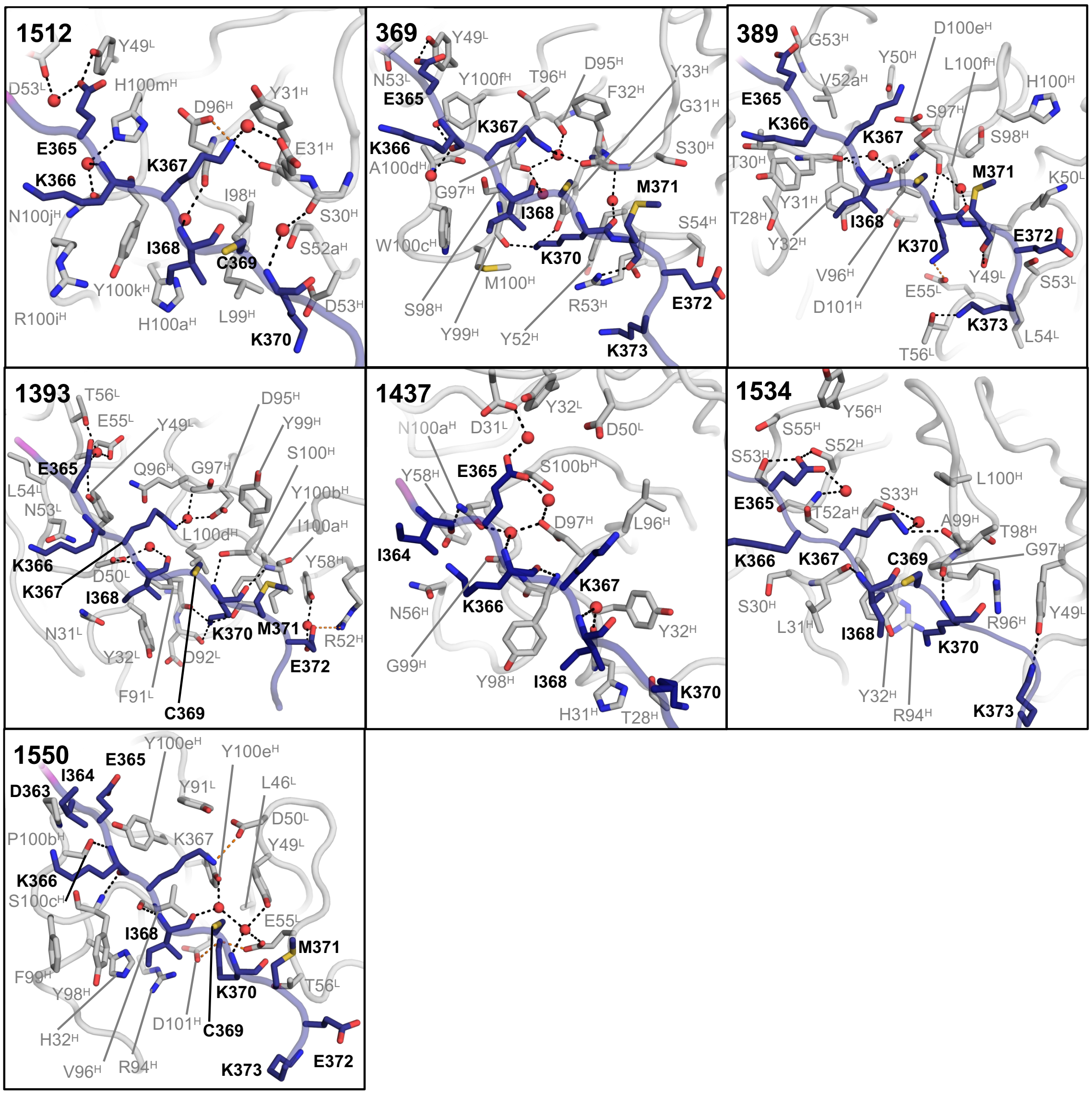


**Figure S3. Molecular interactions of β-epitope binding Fabs with αTSR CS-T3 epitope.** Fab (gray) protein backbone shown in cartoon representation. Residues with more than 5Å$^2$ buried surface area at the Fab:CS-T3 interface, residues forming water-mediated interactions, and relevant backbone segments, are shown as sticks. Waters mediating Fab:CS-T3 binding are shown as red spheres, with hydrogen bond (black) and salt-bridge (orange) interactions as dashed lines. Previously characterized Fab 1512 (PDB:7RXP) is shown for reference.

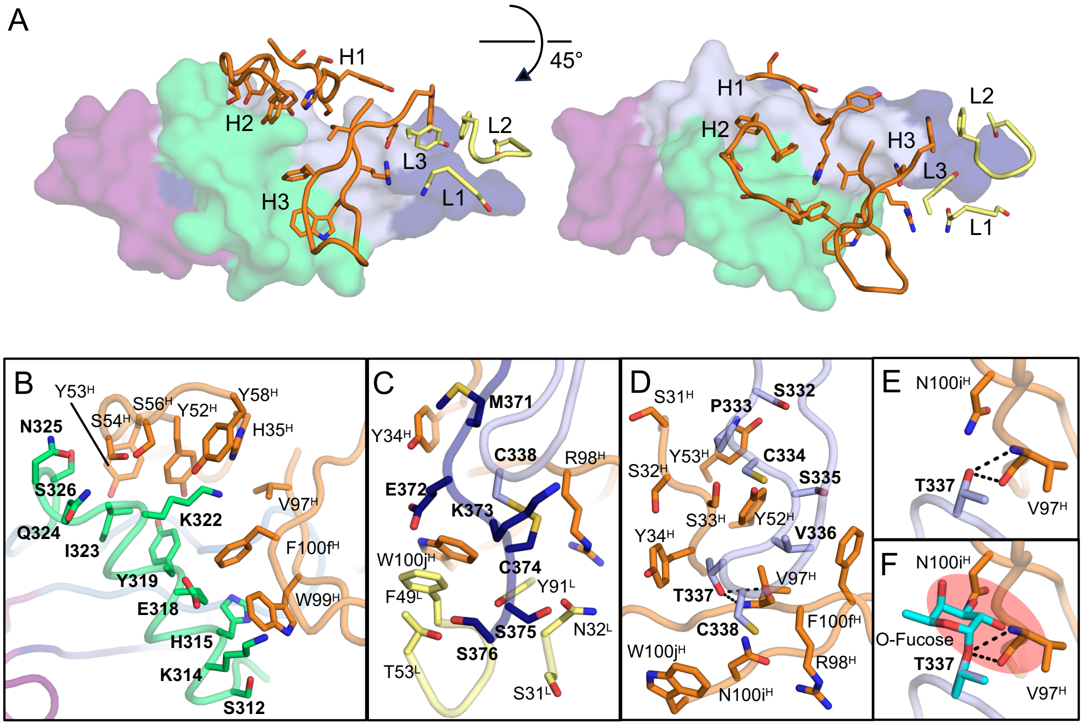


**Figure S4. Paratopes and molecular interactions of Fab 1504 bound to *Pf*CSP αTSR domain.** A) αTSR surface with Fab 1504 paratope residues forming >5Å$^2$ BSA shown as sticks and CDRs labelled H1–3 and L1–3. Molecular interactions at the interface of Fab 1504 with B) Th2R epitope, C) CS-T3 epitope, and D) RII$^+$ epitope. E) Hydrogen bonding interactions of Thr337. F) Thr337 modeled with an O-fucosylated threonine showing potential steric clashes in a red oval. Fab heavy chain (orange) and light chain (yellow) backbone are shown as tubes with side chains in sticks. The αTSR domain is shown with Th2R in green, Th3R in magenta, TSR-homology domain and RII$^+$ in light blue, and CS-T3 in dark blue.

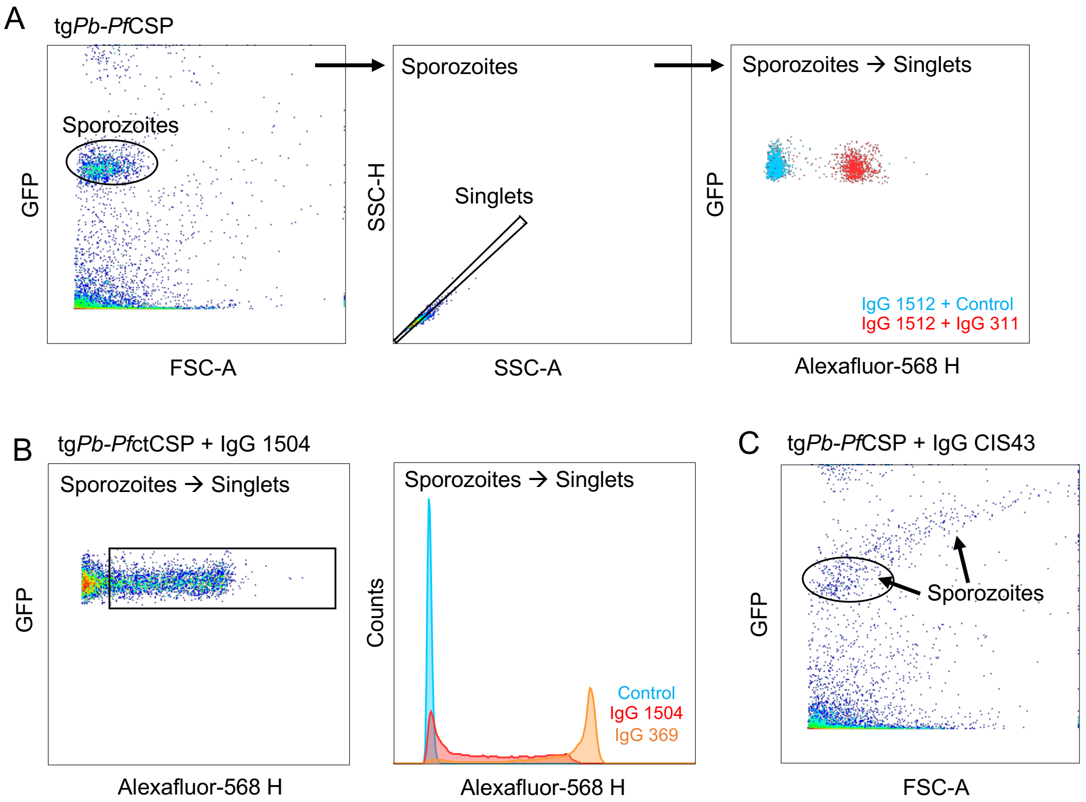


**Figure S5. Antibody binding to transgenic sporozoites by flow cytometry.** A) Representative flow cytometry pseudo-colored plot depicting the gating strategy used to measure antibody binding in all experiments. B) Representative pseudo-colored plot showing binding of IgG 1504 to tg*Pb-Pf*ctCSP sporozoites and a histogram overlaying the binding of IgG 369, IgG 1504, and negative control. C) Flow-cytometry pseudo-colored plot of tg*Pb-Pf*CSP sporozoites bound to IgG CIS43 showing a smeared population of sporozoites marked by black arrows, indicative of morphological changes associated with CSP shedding.

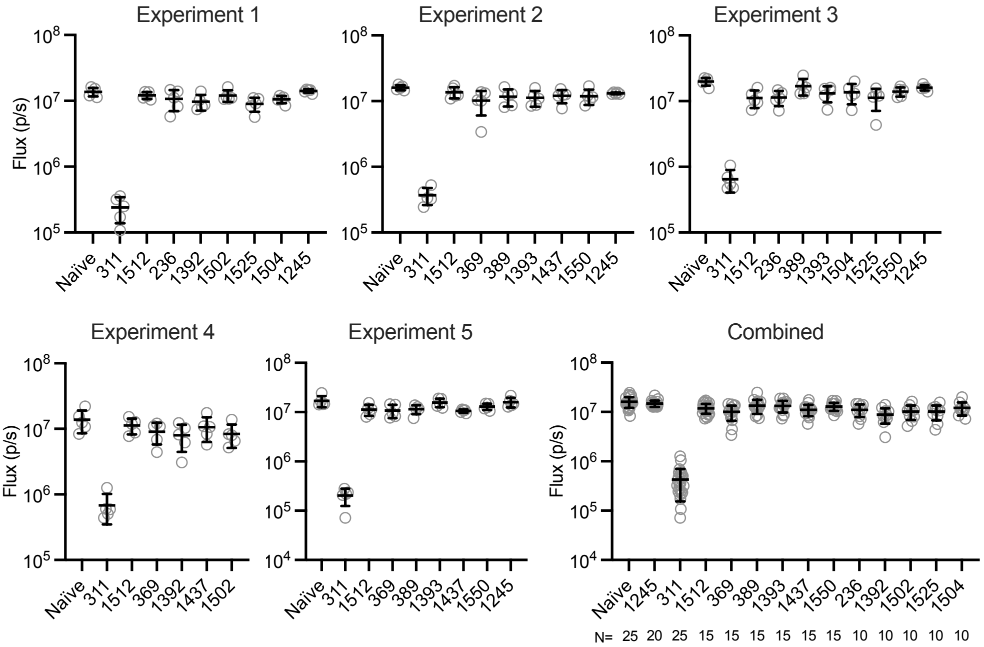


**Figure S6. MAL071 mAbs are weakly protective from *in vivo* sporozoite challenge in mice.** Female C57Bl/6 mice were injected intravenously with IgG1 (300 µg/mouse) 16 hours prior to sporozoite challenge with 2000 transgenic *P. berghei* sporozoites expressing *Pf*CSP and luciferase. Forty-two hours post-challenge, mice were injected with D-luciferin and the liver sporozoite burden measured as flux.

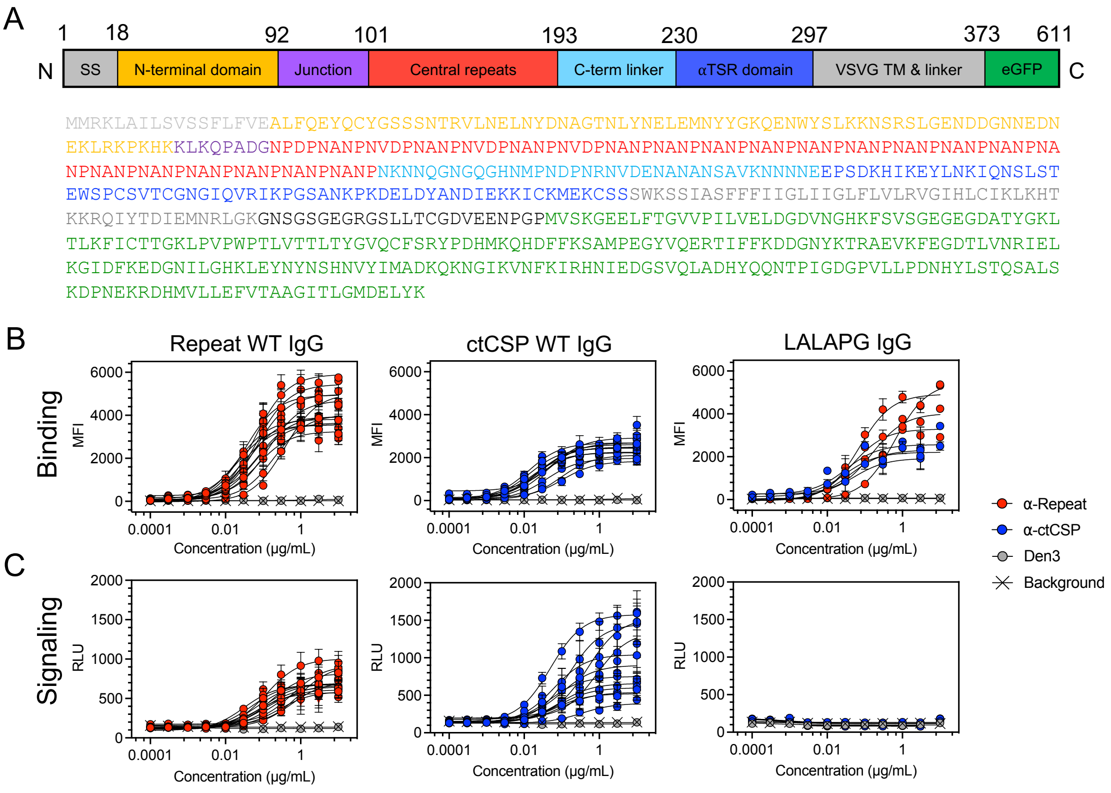


**Figure S7. IgG bound to cell-surface short CSP signal through FcɣRIIIa.** A) Sequence and domain architecture schematic of short CSP amplicon cloned into pLENTI vector for HEK293T cell surface expression. B) Binding and C) FcɣRIIIa signal of CSP mAbs, with three parameter logistic (3PL) curves.

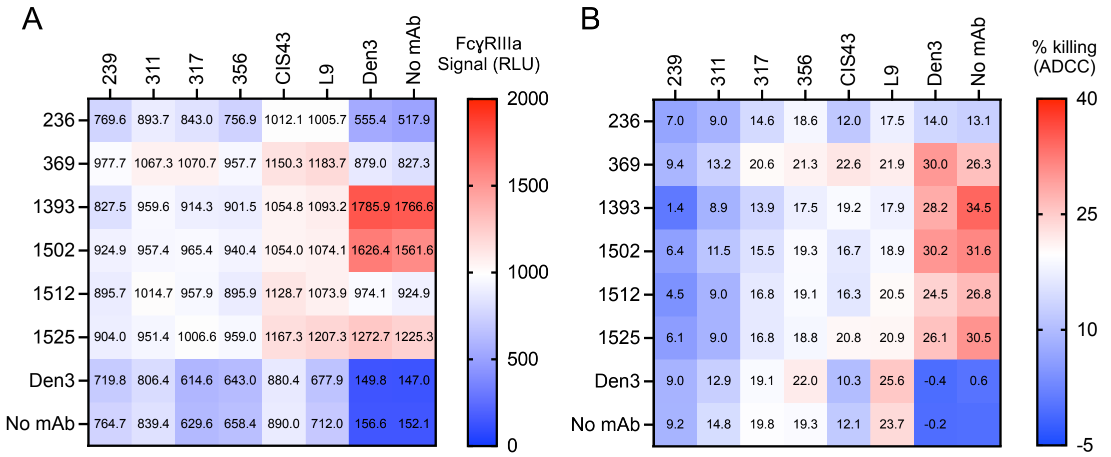


**Figure S8. FcɣRIIIa signaling and ADCC of PfCSP mAbs in combination.** Target cells expressing cell-surface sCSP were incubated with combination of mAbs at 3 µg/ml and added to FcɣRIIIa reporter cells or primary human NK cells. A) FcɣRIIIa signal in Jurkat-Lucia NFAT-CD16 reporter cells and B) percent killing (ADCC) of target cells by primary NK cells.

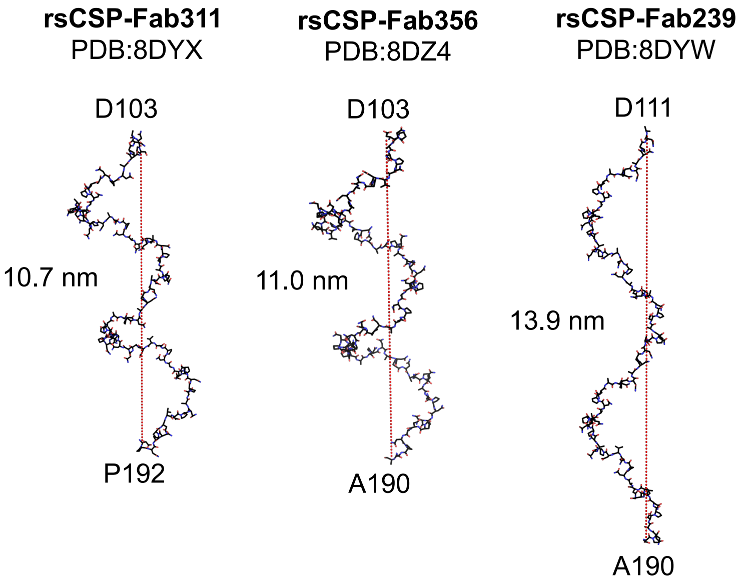


**Figure S9. CryoEM structures and length of rsCSP central repeat region bound to $V_H$3-33 Fabs.** rsCSP bound by Fab shown as sticks, with distance between terminal residues in the EM structures labeled (in nm) and marked by a dotted line. These rsCSP conformations were taken from previous published structures as indicated from their PDB code.

### Supplementary tables

**Table S1.** Characteristics of ctCSP and repeat mAbs used in this study.

| mAb ctCSP | $V_H$ | $V_H$% identity (nt) | $V_H$ SHM (aa) | HCDR3 (aa) | HCDR3 aa sequence | $V_L/_K$ | $V_L$ % identity (nt) | $V_L$ SHM (aa) | LCDR3 (aa) | LCDR3 aa sequence |
|---|---|---|---|---|---|---|---|---|---|---|
| **234** | 3-21*03 | 72 | 8 | 11 | ARGFIQFHYYM | L3-21*03 | 71 | 6 | 11 | QVWHSSSDPVV |
| **236** | 3-48*01 | 70 | 9 | 13 | GGSLHPSAGADWI | L3-1*01 | 72 | 8 | 9 | QAWDSNTYV |
| **352*** | 3-21*04 | 73 | 11 | 9 | GMGIAVRRF | L3-21*02 | 75 | 6 | 10 | QVWDSSTVAS |
| **367*** | 3-21*03 | 74 | 11 | 13 | DGNVMIRGTGDWF | L3-1*01 | 74 | 7 | 10 | QTWDSSTLWV |
| **369** | 4-4*09 | 72 | 12 | 14 | DTGSYMKGWGDYGM | L1-40*01 | 76 | 3 | 10 | QSYDSSLYVV |
| **389** | 3-48*04 | 68 | 15 | 13 | GVSSGHYGTEDLL | K1-5*01 | 70 | 7 | 10 | HQYSSYSPRS |
| **1392** | 4-59*01 | 75 | 19 | 17 | ARDYYDFLTRTYRNFYF | K1-33*01 | 74 | 9 | 8 | QHYDLYPL |
| **1393** | 3-23*01 | 79 | 6 | 12 | DQGGYSIYPLGM | K1-33*01 | 77 | 3 | 9 | QQFDNLPLT |
| **1437** | 3-23*04 | 79 | 8 | 11 | SLDYGGNSLSP | L1-51*01 | 79 | 5 | 11 | GTWDSSLSAGV |
| **1488** | 3-21*04 | 79 | 5 | 10 | RAGGFDAYYF | L3-1*01 | 82 | 2 | 9 | QAWDSSTVV |
| **1502** | 3-21*04 | 77 | 11 | 12 | IFGIASADTTF | L3-21*01 | 79 | 7 | 9 | QVWDSYTAV |
| **1504*** | 4-30-4*10 | 77 | 10 | 17 | LGVRWELLVGGFVNNWF | K3D-15*01 | 78 | 2 | 9 | QQYNNWLGT |
| **1512** | 3-30-3*01 | 80 | 10 | 21 | DDNILGHCRSKSCQRNYYHGM | K2-28*01 | 72 | 3 | 9 | MQALQTPFT |
| **1525** | 3-11*04 | 79 | 10 | 13 | GGGYPQLTHSYYM | L3-1*01 | 80 | 8 | 9 | QAWDSSAVV |
| **1534*** | 3-21*04 | 79 | 6 | 15 | DRGTALRNFEWSDAS | K3-20*01 | 76 | 4 | 8 | QQYGNSHT |
| **1550** | 3-23*01 | 78 | 7 | 13 | DVSYFDSPGYYYF | K1-33*01 | 77 | 4 | 9 | QQYDNLPLT |

| mAb repeat | $V_H$ | $V_H$% identity (nt) | $V_H$ SHM (aa) | HCDR3 (aa) | HCDR3 aa sequence | $V_L/_K$ | $V_L$ % identity (nt) | $V_L$ SHM (aa) | LCDR3 (aa) | LCDR3 aa sequence |
|---|---|---|---|---|---|---|---|---|---|---|
| **224** | 3-49*04 | 93 | 12 | 16 | VQLDYGPGYQYYGMDV | L1-40*01 | 97 | 7 | 11 | QSYDTSLNGWA |
| **227** | 3-33*01 | 94 | 10 | 15 | VKNYESSGYSQCLDY | L1-40*01 | 98 | 2 | 11 | QSYDSSLSAFV |
| **239** | 3-33*06 | 95 | 11 | 12 | DWGGASDRVFDY | K1-5*03 | 94 | 10 | 10 | QHYNSYSRIT |
| **311** | 3-33*01 | 95 | 10 | 12 | AAYYDTSGYGDY | L1-40*01 | 97 | 5 | 12 | QSYDRRLSGSWV |
| **317** | 3-30*04 | 95 | 10 | 10 | DGYSSSFFDF | K1-5*03 | 95 | 8 | 9 | QHYNSYFVT |
| **337** | 3-33*01 | 94 | 12 | 15 | DGADYFDSTGGAFDI | K3-15*01 | 97 | 3 | 8 | QQYKTWWT |
| **356** | 3-33*01 | 94 | 11 | 15 | DSLFYDHDNSGYYGY | K3-15*01 | 98 | 5 | 8 | QQYNNGFT |
| **364** | 3-33*06 | 95 | 10 | 10 | VHDDEPTQDY | K1-5*03 | 95 | 7 | 8 | QQYKRYWT |
| **397** | 3-15*01 | 94 | 10 | 10 | DRDFYRSGGS | K2-28*01 | 98 | 5 | 9 | MQTLQTPHT |
| **399** | 3-49*03 | 95 | 9 | 10 | VGVVIATAVY | K2D-29*02 | 96 | 7 | 9 | MQRIDLPWT |
| **7088** | 3-33*01 | 93 | 13 | 14 | VRFSVGPHGSAFDL | K1-5*01 | 98 | 1 | 10 | QQYNSYSFWT |
| **7150** | 3-15*01 | 95 | 11 | 10 | DRDFYRSGGY | K2-28*01 | 97 | 5 | 9 | MQSLQTPHT |
| **L9** | 3-33*03 | - | 9 | 10 | NFYDGSGPFDY | K1-5*05 | - | 12 | 9 | QEYTSYGRT |
| **CIS43** | 1-3*01 | - | 6 | 12 | LTVLTPDDAFDI | K4-1*02 | - | 3 | 9 | HQYYSSPLT |

*mAb not used in FcγRIIIa reporter assays

**Table S2.** Crystallographic data collection and refinement

| **Data collection** | **Fab367-αTSR** | **Fab369-αTSR** | **Fab389-αTSR** | **Fab1392-αTSR** |
|---|---|---|---|---|
| Beamline | SSRL 12-1 | ALS 501 | APS 23-ID-D | APS 23ID-B |
| Wavelength (Å) | 0.97946 | 0.97741 | 1.0332 | 1.0332 |
| Space group | $P\ 2_1\ 2_1\ 2_1$ | C 2 | C 2 | $C\ 2\ 2\ 2_1$ |
| Cell dimensions | | | | |
| a, b, c (Å) | 40.9, 69.5, 240.1 | 78.3, 71.1, 102.0 | 195.5, 63.7, 139.7 | 95.4, 150.1, 136.4 |
| α, β, ɣ (°) | 90, 90, 90 | 90, 102.8, 90 | 90, 101.8, 90 | 90, 90, 90 |
| Resolution (Å) | 50.0-2.30 (2.34-2.30) | 49.7-2.13 (2.19-2.13) | 48.3-1.75 (1.78-1.75) | 42.1-1.63 (1.66-1.63) |
| $R_{merge}$ | 0.104 (1.36) | 0.207 (1.53) | 0.118 (1.88) | 0.083 (1.26) |
| $R_{meas}$ | 0.120 (1.57) | 0.225 (1.71) | 0.129 (2.08) | 0.086 (1.41) |
| $R_{pim}$ | 0.058 (0.77) | 0.087 (0.74) | 0.052 (0.88) | 0.024 (0.61) |
| Unique reflections | 28,541 (1281) | 30,592 (2456) | 166,880 (8265) | 115,516 (3858) |
| $CC_{1/2}$ | 0.99 (0.39) | 0.99 (0.32) | 0.99 (0.34) | 1.00 (0.42) |
| I / sI | 10.2 (1.2) | 8.2 (1.0) | 11.5 (0.8) | 25.5 (0.9) |
| Completeness (%) | 91.9 (85.5) | 99.7 (96.8) | 99.9 (99.9) | 95.3 (64.2) |
| Redundancy | 3.8 (3.6) | 6.6 (5.0) | 5.8 (5.5) | 11.1 (4.4) |
| **Refinement** | | | | |
| Resolution (Å) | 35.2-2.31 | 49.3-2.13 | 45.6-1.75 | 42.1-1.63 |
| No. reflections | 28,425 | 30,588 | 166,016 | 114,853 |
| Reflections in $R_{free}$ | 1456 | 1521 | 8212 | 5617 |
| $R_{work}$ / $R_{free}$ | 0.24/0.28 | 0.19/0.23 | 0.19/0.21 | 0.17/0.19 |
| No. atoms | 3798 | 4117 | 16136 | 8363 |
| Protein | 3762 | 3808 | 15515 | 7825 |
| Ligand/ion | 0 | 0 | 89 | 341 |
| Water | 36 | 309 | 621 | 197 |
| **B-values (Å$^2$)** | | | | |
| Average B | 65 | 34 | 39 | 38 |
| Fab | 60 | 29 | 39 | 36 |
| αTSR domain | 107 | 59 | 41 | 40 |
| Ligand/ion | 0 | 0 | 50 | 58 |
| Water | 41 | 38 | 40 | 42 |
| Wilson B-value | 42 | 29 | 29 | 28 |
| **R.m.s. deviations** | | | | |
| Bond lengths (Å) | 0.003 | 0.005 | 0.019 | 0.017 |
| Bond angles (°) | 0.58 | 0.73 | 1.47 | 1.48 |
| **Ramachandran Plot(%)** | | | | |
| Favored | 95.8 | 97.2 | 97.8 | 98.0 |
| Allowed | 4.0 | 2.8 | 2.2 | 2.0 |
| Outliers | 0.2 | 0.0 | 0.0 | 0.0 |
| Clashscore | 5.8 | 2.9 | 1.8 | 2.6 |
| **PDB accession code** | 9O8O | 9NI2 | 9NI1 | 9NCY |

Numbers in parentheses refer to the highest resolution shell.

$R_{merge} = \Sigma_{hkl} \Sigma_i \mid I_{hkl,i} - \langle I_{hkl} \rangle \mid / \Sigma_{hkl} \Sigma_i I_{hkl,i}$

$R_{meas} = \Sigma_{hkl} (n/(n-1))^{1/2} \Sigma_i \mid I_{hkl,i} - \langle I_{hkl} \rangle \mid / \Sigma_{hkl} \Sigma_i I_{hkl,I}$

$R_{pim} = \Sigma_{hkl} (1/(n-1))^{1/2} \Sigma_i \mid I_{hkl,i} - \langle I_{hkl} \rangle \mid / \Sigma_{hkl} \Sigma_i I_{hkl,i}$

Where $I_{hkl,i}$ is the scaled intensity of the i$^{th}$ measurement of reflection h, k, l, $\langle I_{hkl} \rangle$ is the average intensity for that reflection, and *n* is the redundancy.

$CC_{1/2}$ = Pearson correlation coefficient between two random half datasets.

$R_{work} = \Sigma_{hkl} \mid F_o - F_c \mid / \Sigma_{hkl} \mid F_o \mid$ x 100, where $F_o$ and $F_c$ are the observed and calculated structure factors, respectively.

$R_{free}$ was calculated as for $R_{work}$, but on a test set comprising 5% of the data excluded from refinement.

Ramachandran values were calculated from MolProbity [1]

**Table S3.** Crystallographic data collection and refinement

| **Data collection** | **Fab1393-αTSR** | **Fab1437-αTSR** | **$F(ab)_2$1502-αTSR** | **Fab1504-αTSR** |
|---|---|---|---|---|
| Beamline | APS 23-ID-B | SSRL 12-2 | ALS 8.2.2 | NSLS-II 17-ID-2 |
| Wavelength (Å) | 1.0332 | 0.97946 | 1 | 0.97934 |
| Space group | $P\ 1\ 2_1\ 1$ | $C\ 2\ 2\ 2_1$ | $I\ 4_1$ | P 1 |
| Cell dimensions | | | | |
| a, b, c (Å) | 71.7, 61.1, 72.1 | 71.7, 106.1, 152.4 | 103.0, 103.0, 172.4 | 72.7, 85.3, 104.3 |
| α, β, γ (°) | 90, 106.2, 90 | 90, 90, 90 | 90, 90 90 | 66.8, 70.3, 65.5 |
| Resolution (Å) | 50.0-1.80 (1.83-1.80) | 50.0-1.95 (1.98-1.95) | 44.5-2.00 (2.03-2.00) | 50.0-2.2 (2.24-2.20) |
| $R_{merge}$ | 0.108 (1.09) | 0.145 (1.68) | 0.146 (3.88) | 0.125 (0.49) |
| $R_{meas}$ | 0.127 (1.30) | 0.159 (1.88) | 0.152 (4.02) | 0.149 (0.62) |
| $R_{pim}$ | 0.068 (0.70) | 0.062 (0.82) | 0.041 (1.07) | 0.079 (0.37) |
| Unique reflections | 55,017 (2670) | 42,265 (2094) | 59,911 (2876) | 98,068 (3510) |
| $CC_{1/2}$ | 0.99 (0.36) | 0.98 (0.42) | 1.00 (0.36) | 0.98 (0.53) |
| I / sI | 12.0 (0.9) | 7.3 (1.1) | 18.7 (0.7) | 8.8 (1.4) |
| Completeness (%) | 98.6 (98.5) | 99.7 (99.8) | 98.9 (94.9) | 94.1 (67.3) |
| Redundancy | 3.4 (3.3) | 6.3 (5.0) | 13.3 (13.5) | 3.3 (2.1) |
| **Refinement** | | | | |
| Resolution (Å) | 43.17-1.80 (1.83-1.80) | 46.85-1.95 (2.00-1.95) | 44.5-2.00 (2.03-2.00) | 46.88-2.19 (2.22-2.19) |
| No. reflections | 54,468 | 42,083 | 59,445 | 97,716 |
| Reflections in $R_{free}$ | 2605 | 2126 | 2974 | 4793 |
| $R_{work}$ / $R_{free}$ | 0.18/0.22 | 0.23/0.26 | 0.21/0.24 | 0.21/0.25 |
| No. atoms | 8932 | 7814 | 7493 | 30937 |
| Protein | 8458 | 7548 | 7303 | 30421 |
| Ligand/ion | 0 | 14 | 48 | 0 |
| Water | 474 | 252 | 142 | 516 |
| **B-values (Å$^2$)** | | | | |
| Average B | 36 | 40 | 52 | 38 |
| Fab | 28 | 38 | 54 | 37 |
| αTSR domain | 56 | 55 | 45 | 46 |
| Ligand/ion | 0 | 35 | 68 | 0 |
| Water | 34 | 34 | 41 | 30 |
| Wilson B-value | 22 | 29 | 35 | 29 |
| **R.m.s. deviations** | | | | |
| Bond lengths (Å) | 0.007 | 0.002 | 0.014 | 0.002 |
| Bond angles (°) | 0.82 | 0.61 | 1.10 | 0.56 |
| **Ramachandran Plots(%)** | | | | |
| Favored | 97.8 | 97.6 | 96.9 | 96.9 |
| Allowed | 2.2 | 2.4 | 2.9 | 2.8 |
| Outliers | 0.0 | 0.0 | 0.2 | 0.3 |
| Clashscore | 1 | 0.8 | 2.6 | 3.4 |
| **PDB accession code** | 9NIY | 9NJ4 | 9NHY | 9O18 |

Numbers in parentheses refer to the highest resolution shell.

$R_{merge} = \Sigma_{hkl} \Sigma_i \mid I_{hkl,i} - \langle I_{hkl} \rangle \mid / \Sigma_{hkl} \Sigma_i I_{hkl,i}$

$R_{meas} = \Sigma_{hkl} (n/(n-1))^{1/2} \Sigma_i \mid I_{hkl,i} - \langle I_{hkl} \rangle \mid / \Sigma_{hkl} \Sigma_i I_{hkl,I}$

$R_{pim} = \Sigma_{hkl} (1/(n-1))^{1/2} \Sigma_i \mid I_{hkl,i} - \langle I_{hkl} \rangle \mid / \Sigma_{hkl} \Sigma_i I_{hkl,i}$

Where $I_{hkl,i}$ is the scaled intensity of the $i^{th}$ measurement of reflection h, k, l, $\langle I_{hkl} \rangle$ is the average intensity for that reflection, and $n$ is the redundancy.

$CC_{1/2}$ = Pearson correlation coefficient between two random half datasets.

$R_{work} = \Sigma_{hkl} \mid F_o - F_c \mid / \Sigma_{hkl} \mid F_o \mid$ x 100, where $F_o$ and $F_c$ are the observed and calculated structure factors, respectively.

$R_{free}$ was calculated as for $R_{work}$, but on a test set comprising 5% of the data excluded from refinement.

Ramachandran values were calculated from MolProbity [1]

**Table S4.** Crystallographic data collection and refinement

| **Data collection** | **Fab1525-αTSR** | **Fab1534-αTSR** | **Fab1550-αTSR** |
|---|---|---|---|
| Beamline | APS 23-ID-B | SSRL 12-1 | SSRL 12-1 |
| Wavelength (Å) | 1.0332 | 0.97946 | 0.97946 |
| Space group | $P\ 6_1\ 2\ 2$ | $P\ 2_12_12_1$ | C 2 |
| Cell dimensions | | | |
| a, b, c (Å) | 186.3, 186.3, 68.6 | 75.8, 86.8, 185.4 | 147.3, 61.0, 70.4 |
| α, β, ɣ (°) | 90, 90, 120 | 90, 90, 90 | 90, 106.5, 90 |
| Resolution (Å) | 46.6-2.20 (2.24-2.20) | 50.0-2.10 (2.14-2.10) | 50.0-2.0 (2.03-2.00) |
| $R_{merge}$ | 0.071 (0.42) | 0.202 (1.43) | 0.130 (0.35) |
| $R_{meas}$ | 0.072 (0.44) | 0.222 (1.58) | 0.145 (0.42) |
| $R_{pim}$ | 0.012 (0.12) | 0.090 (0.65) | 0.061 (0.23) |
| Unique reflections | 31,697 (912) | 65,858 (2964) | 36,750 (1081) |
| $CC_{1/2}$ | 1.00 (0.94) | 0.99 (0.43) | 0.98 (0.84) |
| I / sI | 53.0 (6.3) | 8.1 (1.0) | 11.5 (3.9) |
| Completeness (%) | 87.7 (51.7) | 96.6 (88.1) | 90.2 (52.7) |
| Redundancy | 33.7 (13.9) | 5.7 (5.2) | 5.3 (2.8) |
| **Refinement** | | | |
| Resolution (Å) | 46.6-2.20 (2.24-2.20) | 42.5-2.09 (2.12-2.09) | 37.27-2.00 (2.03-2.00) |
| No. reflections | 31,478 | 65,634 | 36,710 |
| Reflections in $R_{free}$ | 1576 | 3295 | 1859 |
| $R_{work}$ / $R_{free}$ | 0.20/0.24 | 0.21/0.25 | 0.17/0.21 |
| No. atoms | 8471 | 15647 | 8642 |
| Protein | 8189 | 15202 | 8366 |
| Ligand/ion | 56 | 0 | 0 |
| Water | 226 | 445 | 276 |
| **B-values (Å$^2$)** | | | |
| Average B | 56 | 36 | 30 |
| Fab | 53 | 34 | 26 |
| αTSR domain | 53 | 53 | 44 |
| Ligand/ion | 66 | 0 | 0 |
| Water | 44 | 32 | 30 |
| Wilson B-value | 39 | 25 | 21 |
| **R.m.s. deviations** | | | |
| Bond lengths (Å) | 0.003 | 0.003 | 0.01 |
| Bond angles (°) | 0.56 | 0.58 | 0.82 |
| **Ramachandran Plots(%)** | | | |
| Favored | 97.6 | 97.1 | 97.6 |
| Allowed | 2.2 | 2.9 | 2.4 |
| Outliers | 0.2 | 0.0 | 0.0 |
| Clashscore | 1.9 | 1.6 | 1.9 |
| **PDB accession code** | 9NDM | 9NHV | 9NIW |

Numbers in parentheses refer to the highest resolution shell.

$R_{merge} = \Sigma_{hkl} \Sigma_i \mid I_{hkl,i} - \langle I_{hkl} \rangle \mid / \Sigma_{hkl} \Sigma_i I_{hkl,i}$

$R_{meas} = \Sigma_{hkl} (n/(n-1))^{1/2} \Sigma_i \mid I_{hkl,i} - \langle I_{hkl} \rangle \mid / \Sigma_{hkl} \Sigma_i I_{hkl,I}$

$R_{pim} = \Sigma_{hkl} (1/(n-1))^{1/2} \Sigma_i \mid I_{hkl,i} - \langle I_{hkl} \rangle \mid / \Sigma_{hkl} \Sigma_i I_{hkl,i}$

Where $I_{hkl,i}$ is the scaled intensity of the $i^{th}$ measurement of reflection h, k, l, $\langle I_{hkl} \rangle$ is the average intensity for that reflection, and *n* is the redundancy.

$CC_{1/2}$ = Pearson correlation coefficient between two random half datasets.

$R_{work} = \Sigma_{hkl} \mid F_o - F_c \mid / \Sigma_{hkl} \mid F_o \mid$ x 100, where $F_o$ and $F_c$ are the observed and calculated structure factors, respectively.

$R_{free}$ was calculated as for $R_{work}$, but on a test set comprising 5% of the data excluded from refinement.

Ramachandran values were calculated from MolProbity [1]

**Table S5.** Hydrophobic HCDR3 residues (highlighted) buried in αTSR core.

| mAb | AA | BSA (Å$^2$) | % BSA | HCDR3 sequence |
|---|---|---|---|---|
| 234 | $F98^H$, $F100a^H$ | 162 | 23 | ARGFIQFHYYM |
| 236 | $L98^H$ | 76 | 13 | GGSLHPSAGADWI |
| 352 | $I98^H$ | 100 | 15 | GMGIAVRRF |
| 367 | $I100^H$ | 72 | 10 | DGNVMIRGTGDWF |
| 1392 | $F100a^H$ | 117 | 15 | ARDYYDFLTRTYRNFYF |
| 1488 | $F99^H$ | 122 | 21 | RAGGFDAYYF |
| 1502 | $I98^H$ | 91 | 14 | IFGIASADTTF |
| 1525 | $L100a^H$ | 101 | 15 | GGGYPQLTHSYYM |

**Table S6.** Fab-αTSR crystallization and cryoprotection conditions

| Complex | Crystallization conditions | Cryoprotectant |
|---|---|---|
| Fab367-αTSR | 40% PEG-400<br>0.1 M acetate pH 4.5, final pH 5.4 | 30% glycerol |
| Fab369-αTSR | 0.17 M ammonium sulphate<br>15% glycerol<br>25.5% PEG-4000 | 30% glycerol |
| Fab389-αTSR | 2.0 M ammonium sulphate<br>0.2 M lithium sulphate<br>0.1 M Cyclohexyl-3-aminopropane sulfonic acid pH 10.5 | 30% glycerol |
| Fab1392-αTSR | 30% PEG-600<br>10% (v/v) glycerol<br>0.5 M ammonium sulfate<br>0.1M Tris pH 7.0 final pH 5.6 | 30% glycerol |
| Fab1393-αTSR | 20% PEG 3350<br>0.2 M di-ammonium tartrate pH 6.6<br>Protein G domain III | 30% ethylene glycol |
| Fab1437-αTSR | 8.5% isopropanol<br>17% PEG-4000<br>15% glycerol<br>0.085 M HEPES pH 7.5 | 30% ethylene glycol |
| F(ab´)$_2$1502-αTSR | 0.2 M sodium citrate<br>30% (v/v) PEG-400<br>0.1 M Tris pH 8.5 | 30% ethylene glycol |
| Fab1504-αTSR | 36% (v/v) ethanol<br>0.1 M phosphate-citrate pH 4.37<br>5% PEG-1000 | 30% 2-methyl-2,4-pentandiol |
| Fab1525-αTSR | 20% PEG 3350<br>0.2 M Mg-sulfate pH 5.9<br>Protein G domain III | 30% ethylene glycol |
| Fab1534-αTSR | 20% PEG 3350<br>0.2 M K-iodide pH 6.8 | 30% ethylene glycol |
| Fab1550-αTSR | 20% PEG-6000<br>1.0 M Li-chloride<br>0.1 M citric acid pH 4.0<br>Protein G domain III | 30% glycerol |
